# Proximity labeling reveals ZFP36L1 as a central hub for post-transcriptional regulation networks in T cells

**DOI:** 10.1101/2025.07.25.666749

**Authors:** Anouk P. Jurgens, Branka Popović, Floris P.J. van Alphen, Leyma Wardak, Antonia Bradarić, Sander Engels, Carmen van der Zwaan, Maartje van den Biggelaar, Arie J. Hoogendijk, Julien Béthune, Monika C. Wolkers

**Affiliations:** Sanquin Blood Supply Foundation, Department of Research, T cell differentiation lab, Plesmanlaan 125, 1066 CX Amsterdam, The Netherlands; Sanquin Blood Supply Foundation, Department of Research, Bleeding & Hemostasis, Plesmanlaan 125, Amsterdam, The Netherlands; Amsterdam UMC, University of Amsterdam, Landsteiner Laboratory, Amsterdam Institute for Infection & Immunity and Cancer Center Amsterdam, Meibergdreef 9, 1105 AZ Amsterdam, The Netherlands; Oncode Institute, Utrecht, The Netherlands; Department of Biotechnology, HAW Hamburg, Germany

**Keywords:** UltraID, T cell, RNA binding protein, post-transcriptional regulation

## Abstract

Effective T cell responses against pathogens require a rapid yet tightly controlled remodeling of the proteome, and RNA binding proteins (RBPs) are key in this process. For instance, the RBP ZFP36L1 prevents excessive protein production and thereby limits immunopathology. ZFP36L1 is primarily known to mediate mRNA decay, but it can also regulate other processes. How its mode of action relates to its interaction partners is, however, not well-understood. Here, we mapped the ZFP36L1 interactome in primary human T cells. Using proximity labeling, we identified known and new interactors that regulate 3’UTR-mediated RNA degradation, deadenylation, stress granule/p-body formation, as well as 5’UTR-mediated translation repression and mRNA decapping. Snapshot analysis uncovered the ZFP36L1 interactome dynamics and RNA (in)dependency throughout T cell activation. Intriguingly, proximity labeling also uncovered regulators of ZFP36L1 protein expression: This included the helicase UPF1, which not only interacts with ZFP36L1 protein but also promotes its protein expression. Altogether, this comprehensive interactome map underlines the versatility of interactions with ZFP36L1 and their possible role in cellular function.

## INTRODUCTION

T cells play a crucial role in clearing infected and malignant cells from our body. To exert their effector function, T cells rapidly remodel their proteome upon antigen recognition ^1–3^. This includes their ability to produce effector molecules such as granzymes and proinflammatory cytokines, a central feature for their capacity to clear target cells ^4,5^. Yet, tight regulation of gene expression is key for preventing excessive responses, as it can lead to immunopathology ^6,7^. To achieve effective yet balanced immune responses, several regulatory layers are in place, and RNA binding proteins (RBPs) are key contributors herein ^8–11^. Defined by the signals that T cells receive, RBPs control the magnitude and duration of cytokine production ^12–14^, and they regulate cell cycle and checkpoint molecule production levels ^15–17^. Notably, mutations in RBPs are linked to autoimmunity in humans ^18,19^, highlighting the significance of this class of proteins in regulating adequate immune responses.

An important RBP for T cell effector function is ZFP36L1. ZFP36L1 belongs to the ZFP36 protein family and mediates mRNA decay of transcripts that contain AU-rich elements (ARE) in their 3’UTR ^20^, as exemplified for mRNA coding for cell cycle proteins and cytokines ^21–25^. ZFP36L1 deletion in T cells results in increased cytokine production, which in turn can restore function in tumor-infiltrating T cells and improve anti-tumor responses in mice ^24^. Intriguingly, ZFP36L1 regulates cytokine expression upon T cell activation in a time-dependent manner ^17,23,24^: in the first 4-24h, ZFP36L1 restrains cytokine production, and it does so at least in part by recruiting the CCR4/CNOT complex ^24^. However, ZFP36L1 can also exert several other functions: together with its sister protein ZFP36L2, ZFP36L1 binds to ready-to-deploy cytokine mRNAs in memory T cells, thereby conferring a block of translation and preventing aberrant production in the absence of T cell stimuli ^26^. Furthermore, ZFP36L1 has been implicated in other cell types in the formation of stress-granules, P-bodies and membraneless structures ^20,27,28^. Provided that most RBPs are highly interactive with RNA and proteins, forming multiple ribonucleoprotein (RNP)^29^ complexes, we hypothesized 1) that ZFP36L1 interacts with a broad set of proteins to carry out diverse functions, and 2) that ZFP36L1 may alter its interactome over the course of T cell activation to modulate its activity. However, to date there is only anecdotal evidence of protein interactions with ZFP36L1 in T cells, and a comprehensive map of the ZFP36L1 interactome throughout T cell activation is lacking.

To shed light on how ZFP36L1 executes its multiple functions, and which interaction partners it employs to do so, we performed intracellular proximity labeling in human T cells using UltraID, an engineered protein biotin ligase, which rapidly biotinylates proteins within a range of 10 nm ^30^. We measured the ZFP36L1 interactome in resting T cells and during T cell activation, and we defined the dependency of RNA for ZFP36L1 interactors. This extensive interactome map uncovered known and previously unreported interactors with ZFP36L1 involved in RNA regulation. In addition, we identified known and novel regulators of ZFP36L1 protein levels. In conclusion, our study yields novel insights into the intricate regulation of gene expression by RBPs, as exemplified here for ZFP36L1.

## RESULTS

### Optimizing proximity labeling in primary human T cells

To perform proximity labeling for ZFP36L1, we fused the biotin ligase variant UltraID ^30^ to the N-terminus of ZFP36L1 (UltraID_L1). To identify UltraID_L1 expressing cells, we co-expressed a GFP reporter gene with ZFP36L1 (UltraID_L1_P2A_GFP; **Figure 1A**). UltraID fused to the C-terminus of GFP (UltraID_GFP) and UltraID alone (UltraID only) served as controls (**Figure 1A**). As previously reported ^30^, clear biotinylation profiles were detected in the human fibrosarcoma cell line FLYRD18 after 10 minutes of incubation with 50 μM biotin (**Supplementary Figure 1A**). This was, however, not the case in human T cells. T cells require biotin-dependent carboxylases for fatty acid synthesis and amino acid metabolism, and proliferating T cells absorb five-fold more biotin than resting T cells ^31–33^. Therefore, increasing the biotin levels to up to 500 µM was required to obtain detectable biotinylation profiles in human T cells (**Supplementary Figure 1B**). Moreover, as 10 minutes appeared not sufficient to attain proximity labeling in human T cells, we increased the incubation time to 16h, when T cell activation plateaus, therefore requiring less biotin (**Supplementary Figure 1B, second & third lane: UltraID_only & UltraID_L1**). Because proximity labeling with UltraID_GFP displayed more specific biotinylated protein bands than UltraID protein alone (**Figure 1B, Supplementary Figure 1B**), and because it included a protein of similar size as ZFP36L1 fused to the UltraID enzyme, we continued with UltraID_GFP as control in all downstream analyses.

**Figure 1.**
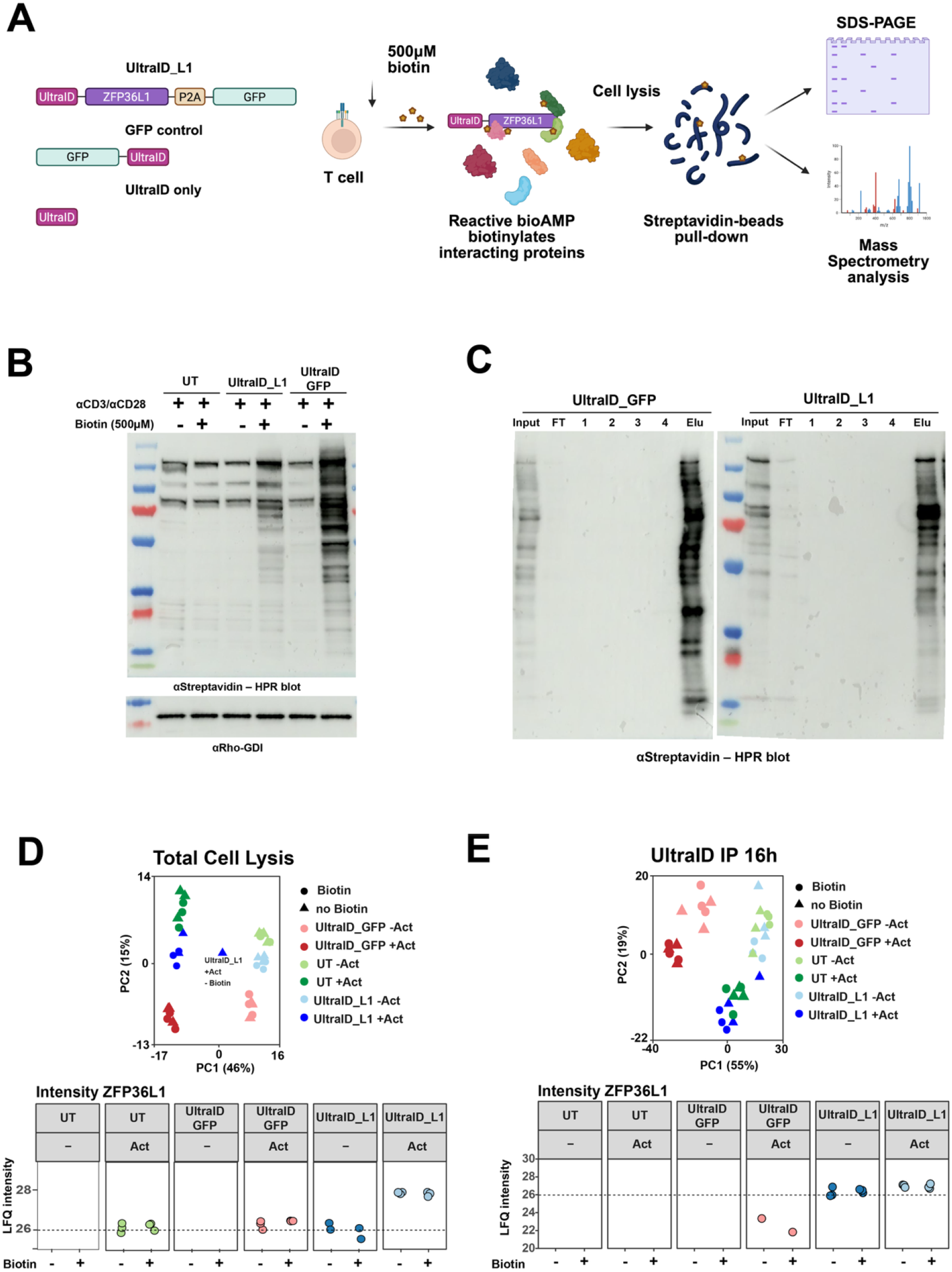
Proximity labeling in primary human T cells. **a**. Schematic representation of proximity labeling protocol in CD3^+^ T cells and UltraID plasmids used. Indicated UltraID BirA* fusion genes were retrovirally transduced into CD3^+^ T cells (n=3 donor pools, 40 donors each). After 6 days, GFP^+^ T cells were FACS-sorted on GFP expression. The next day, T cells were re-activated for 16h with aCD3/aCD28 in the presence or absence of 500 µM biotin. Biotinylated proteins were captured with Streptavidin beads from cell lysates by biotin-streptavidin affinity capture. Biotinylation was confirmed by SDS-PAGE, and biotinylated proteins were identified by Mass Spectrometry. **b**. Immunoblot for biotinylated proteins of untransduced (UT) T cells, and T cells transduced with UltraID_ZFP36L1_GFP (UltraID_L1) or UltraID_GFP (UltraID_GFP). aRho-GDI was used as loading control. Representative blot of two independently performed experiments, 3 donor pools each. **c**. Immunoblot of input (10%), flow through and consecutive washes (1-4) of the pulldown, in addition to the elution (10%). Representative blot of two independently performed experiments, 3 donor pools each. **d**. Top: Principal component analysis (PCA) of total cell lysates. Color indicates constructs expressed in T cells, circle/triangle indicates whether or not biotin was added. Bottom: LFQ intensity of ZFP36L1 in the presence or absence of aCD3/aCD28 activation and biotin. Dotted line indicates endogenous ZFP36L1 levels in untransduced activated T cells. **e**. Top: PCA for biotinylated proteins identified by MS upon streptavidin pulldown in T cell screen 1 of indicated conditions. Bottom: LFQ intensity of ZFP36L1 in the presence or absence of aCD3/aCD28activation and biotin. Dotted line indicates endogenous ZFP36L1 levels in untransduced activated T cells. *See Supplementary Figure 1*

### Proximity labeling identifies the ZFP36L1 interactome in primary human T cells

Having established proximity labeling in human T cells, we questioned which proteins interact with ZFP36L1 upon T cell activation. To avoid alterations in T cell function by exogenous ZFL36L1 expression, we aimed for limited (∼20%) transduction efficiency **(Supplementary Figure 1C)**. This expression level did not influence T cell viability, the expression levels of the activation marker CD69, or the production of TNF, a reported target of ZFP36L1 ^24,34^(**Supplementary Figure 1D, E**). Furthermore, to limit inter-donor variability, we used 3 T cell pools of 40 healthy donors each. FACS-sorted UltraID_L1 and UltraID_GFP expressing T cells were expanded for 7 days and then activated for 16 hours with αCD3/αCD28 in the presence or absence of biotin, a timeframe when ZFP36L1 displays its highest activity in T cells ^17^. Immunoblots revealed distinct biotinylation patterns between UltraID_GFP and UltraID_L1 expressing T cells (**Figure 1C, Supplementary Figure 1F**), indicating differential protein interactions.

Mass spectrometry (MS) analysis of the total cell lysates identified in total 5250 proteins (range 4500-5000 proteins/sample; **Supplementary Figure 2A, Supplementary Table 1)**. Principal component analysis (PCA) confirmed that biotinylation *per se* did not affect the total proteome (**Figure 1D**). Rather, T cell activation was the prime driver of differential protein expression (**Figure 1D, Supplementary Figure 2B**). As expected, differentially expressed proteins between resting and activated T cells included ZFP36L1, which was induced upon T cell activation (**Figure 1D**). MS analysis of isolated biotinylated proteins identified a total of 2439 proteins (range: 1700-2500 proteins/sample; **Supplementary Figure 2C, Supplementary Table 2)**. As expected from its fusion to the proximity labeling enzyme, ZFP36L1 was primarily detected in the UltraID_L1 pull down **(Figure 1E)**. Again, PCA was mainly driven by T cell activation status, together with the expression of UltraID_GFP versus UltraID_L1 (**Figure 1E, Supplementary Figure 2D)**.

To uncover the ZFP36L1 interactome, we utilized supervised classification ^35^. In short, we generated theoretical profiles in which the conditions of interest (interacting with ZFP36L1 in resting T cells (shape 1) or in resting and activated T cells (shape 2)) were set to an arbitrary high and low in all other samples, after which differentially expressed proteins in the ‘high’ sample were correlated with all statistically significant proteins (*for more details see methods section*) **(Figure 2A)**. Using a cut-off of log fold change (LFC) >1, adjusted p-value (p-adj) <0.05, and Pearson correlation ≥0.6, we identified 156 proteins in close proximity to ZFP36L1. 27 ZFP36L1 interactions depended on the presence of UltraID_L1 and were found with T cell activation only (shape 1), and 129 irrespective of the T cell activation status (shape 2; **Figure 2A, B, Supplementary Figure 2E, Supplementary Table 3**). As expected, gene Ontology (GO) analysis from shape 2 identified RNA-related processes, such as mRNA decay, RNP granules, and mRNA binding (**Figure 2C**). Identified proteins included previously reported ZFP36L1 interactors, such as the deadenylating CCR4-CNOT complex members CNOT1, CNOT2, CNOT3, CNOT7, CNOT6L, CNOT10 and CNOT11 (^36–39^**; Figure 2B, C)**. Other known ZFP36L1 interactors were the 14-3-3 protein family members YWHAB and YWHAZ ^40,41^ and the exocyst complex components EXOC1, EXOC3, EXOC4 ^24^ (**Figure 2B)**.

**Figure 2.**
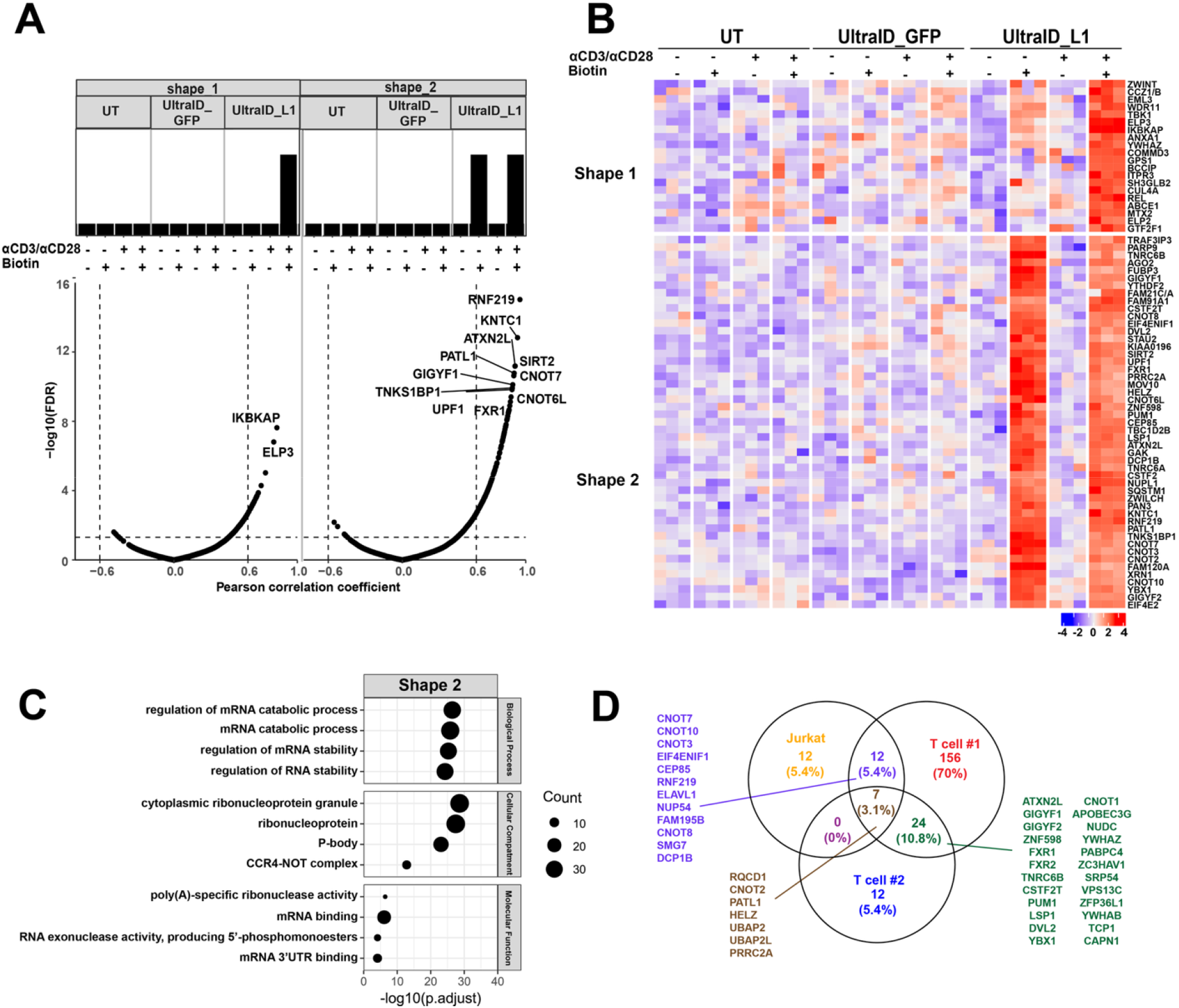
Proximity labeling identifies the ZFP36L1 interactome in primary human T cells. **a**. Supervised classification of biotinylated proteins identified as ZFP36L1 interactors. Top panel indicates which sample(s) were set to an arbitrary high. Shape 1: biotinylated proteins enriched (LFC>1, p.adj <0.05) in activated T cells expressing UltraID_L1 compared to all other conditions. Shape 2: biotinylated proteins enriched in T cells expressing UltraID_L1 irrespective of T cell activation status. Correlation between hits and disorder-specific theoretical protein profiles using a Pearson correlation coefficient of >0.6 (dotted line) and Benjamini-Hochberg adjusted P value (p.adj) <0.05 was used as threshold. **b**. Heatmap of supervised classification displaying biotinylated proteins identified in a). Color scale represents Z-scored log2 median-centered averaged intensities. **c**. GO analysis of identified ZFP36L1 interactors from shape 2. Protein interactors were devided according to biological process, cellular component and molecular function. **d**. Venn diagram depicting the overlap of identified ZFP36L1 interactors (Pearson correlation coefficient >0.5) from shape 1 and 2 for Jurkat cells and two independently performed pulldowns in primary T cells (T cell screen 1 and 2). *See Supplementary Figure 2 & 3*

Repeating the UltraID proximity labeling in primary human T cells confirmed 31 proteins in close proximity to ZFP36L1. Thus, even with the lower coverage of the second screen, UltraID proximity labeling provides robust outcomes (**Figure 2D, Supplementary Figure 3A, Supplementary Table 4, 5)**. Furthermore, an UltraID-L1 pulldown in Jurkat cells yielded similar - yet not identical - proteins in proximity to ZFP36L1 **(Figure 2D, Supplementary Figure 3B-D, Supplementary Table 6, 7**), highlighting the added value of performing proximity labeling in primary cells. Similarly, 32 proteins (6.8%) found in either one of both T cell screens overlapped with a recently published ZFP36L1-APEX proximity labelling in HEK293T cell line, and only 8 proteins were shared between the two T cell screens and HEK293T cells ^42^. Yet, 256 and 183 proteins were specifically enriched in HEK293T cells or T cells, respectively (**Supplementary Figure 3E)**, suggesting that either expression of certain proteins is cell-type specific, or ZFP36L1 has cell-type specific protein interactions, in addition to its core interactome. Combined, we here showcased the potential of UltraID proximity labeling in primary human T cells to measure protein-protein interactions, as exemplified for ZFP36L1.

### ZFP36L1 interactome covers a multitude of RNA-related processes as well as protein turnover

We next zoomed in on the cellular processes annotated for known and candidate interactors of ZFP36L1. Specifically, we examined proteins that were either found in close proximity to ZFP36L1 in at least two out of three screens, or that showed strong correlation with ZFP36L1 proximity in our ‘shapes’ in T cells screen 1 and that were part of a complex with another candidate interactor. As expected, most identified proteins contribute to mRNA decay processes such as mRNA stabilization, de-adenylation, de-capping and degradation (**Figure 3A)**. In addition to the above-mentioned CCR4-NOT complex proteins (**Figure 3A, B**), we identified novel interactions from the de-adenylation complex. This included the PAN2/3 complex proteins **(Figure 3A)**, which trim Poly (A) tails prior to mRNA degradation through the CCR4-Not complex, a rate limiting step of mRNA degradation ^43,44^. We also detected PUM1, which directly binds to the CCR4-NOT complex and supports its function in mRNA de-adenylation ^45^ and TNRC6C, which directly associates with the PAN2/3 and CCR4-NOT complexes ^46^, in addition to its contribution to microRNA degradation (**Figure 3A**). These ubiquitous interactions of ZFP36L1 with most - if not all - proteins involved in mRNA de-adenlyation pathway thus evidence ZFP36L1’s central place in this process. Of note, and in line with previous reports of the cell-cycle dependent nuclear localization of ZFP36L1 ^42,47^, we detected interactions with the nuclear alternative poly-adenylation regulating proteins CSTF2 and CSTF2T ^48^ and with Fam120A, which associates with the splicing protein SRFS1 ^49^ (**Figure 3A**).

**Figure 3.**
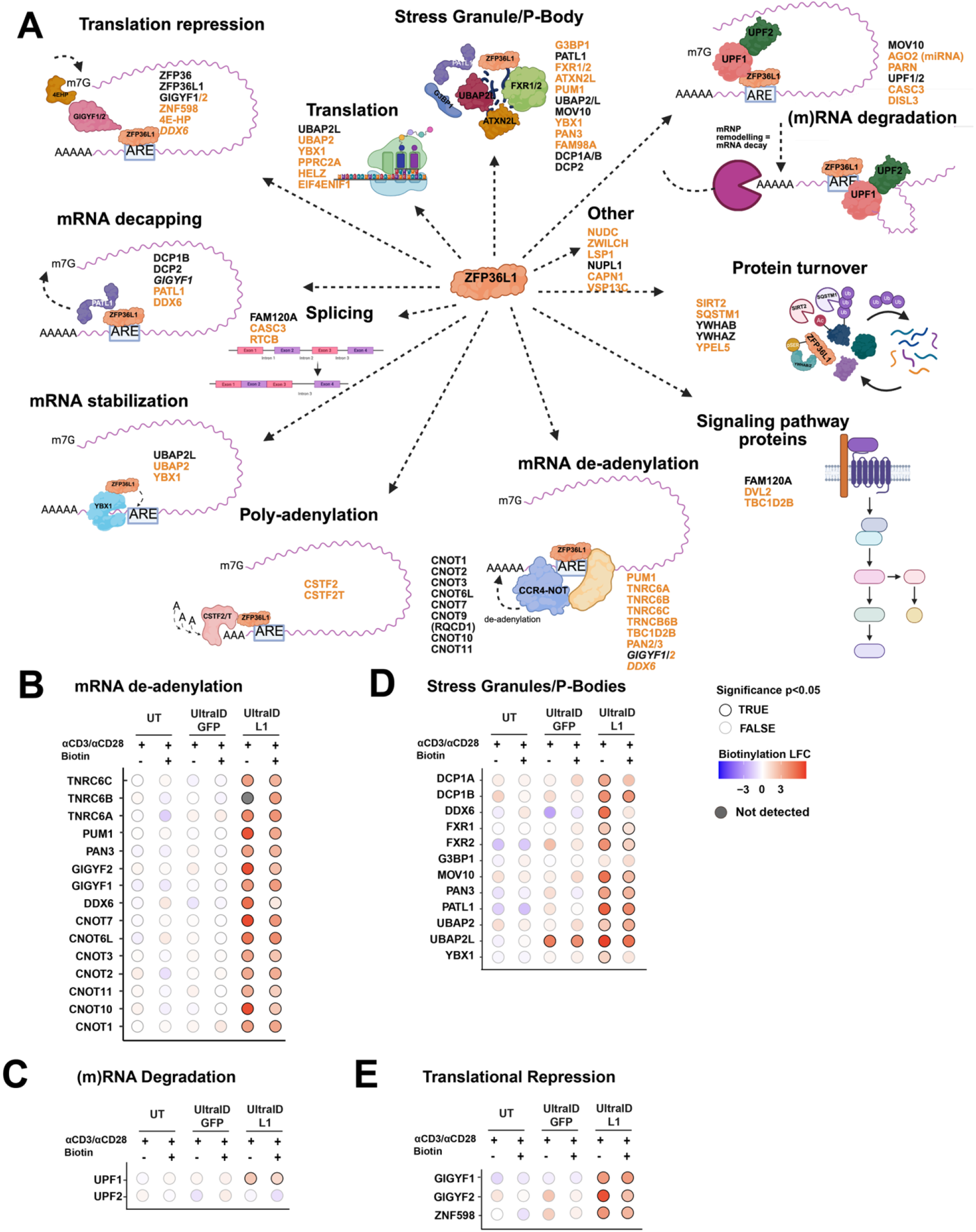
ZFP36L1 interactome correlates with a multitude of RNA control mechanisms. **a**. Network of the ZFP36L1 interactome identified in at least 1/2 proximity labeling T cell screens and/or the Jurkat screen. Cut off for T cell screen 1 was Pearson correlation of 0.6, and for T cell screen 2 and Jurkat screen 0.5, and LFC > 1 and p.adj < 0.05 for all three. Proteins annotated according to molecular function or cellular location. Black: previously reported interactions, Orange: newly identified proteins interacting with ZFP36L1. **b. -e**. LFQ intensity plots of identified ZFP36L1 interactors. Black circles: significantly enriched proteins (p.adj<0.05) in Biotin over no Biotin samples. Grey circles: no significant enrichment. Fold-change (log2FC) enrichment is indicated by color code. n= 3 donor pools.

In the mRNA degradation pathway, we identified the RNA helicase UPF1 as being in close proximity to ZFP36L1, which regulates nonsense-mediated mRNA decay (NMD), Staufen-mediated mRNA decay (SMD), and Regnase-1-mediated mRNA decay ^50–53^ **(Figure 3C)**. ZFP36L1 also interacted with MOV10 (**Figure 3A**), a 5’ to 3’ RNA helicase that supports UPF1 in target mRNA degradation ^54^. Intriguingly, UPF1 deficiency in early murine B cell development resembles the phenotype of ZFP36L1/ZFP36L2 deficiency, which leads to perturbed thymic and B cell development in mice ^52,55^. UPF1 and MOV10 interactions with ZFP36, the sister protein of ZFP36L1, were recently reported in HEK293T cells (Bestehorn et al. 2025) (**Supplementary Figure 2E**), indicating that interactions with UPF1 and MOV10 are conserved in different cell types and between ZFP36 family members. This also holds true for the poly(A) RNase PARN and the RISC complex member AGO2 (**Figure 3A)**, confirming that ZFP36 proteins play a central role in regulating mRNA stability of their target genes.

In line with the described location of ZFP36L1 in stress granules (SG) and P-bodies ^27^, ZFP36L1 interacted with the SG and P-body associated proteins PATL1, UBAP2L, and MOV10 together with G3BP1 (formation of stress granules) and DDX6 (formation of p-bodies) ^56–59^ (**Figure 3A, B, D**). UltraID proximity labeling extended this known list of SG and P-body associated ZFP36L1 candidate interactors to FXR1/2, ATXN2L, PUM1, UBAP2, YBX1, and PAN3 ^43,60,61^.

We also identified proteins in close proximity to ZFP36L1 that are involved in RNA processes regulated through the 5’UTR. This included the decapping proteins DCP2 and DCP1B ^62^ and PATL1, one of the strongest ZFP36L1 candidate interactors (**Figure 3A, D, Supplementary Figure 2E**), which also interacts with DCP1 ^20,43^. DDX6 could also possibly contribute to this 5’UTR interaction, as it can bind to the EDC complex to assist mRNA decapping by DCP1/2 and PATL1 ^63^. Finally, the RBP scaffolding proteins GIGYF1, GIGYF2 and ZNF598, were consistently identified in close proximity to ZFP36L1 **(Figure 3A, E**). GIGYF1 and GIGYF2 interact with DDX6 and with the translation repressor 4EHP (Tollenaere et al. 2019; Weber et al. 2020), both of which we also found interacting with ZFP36L1 (**Figure 3A, 2B)**. Interestingly, ZFP36L1 also interacts with factors that can promote translation, such as UBAP2 and UBAP2L proteins ^64^, as well as the m6A reader PPRC2A, which is reported to regulate mRNA metabolism and facilitate translation of uORF-containing transcripts by promoting leaky scanning ^65^ (**Figure 3A**).

In addition to RBP-ZFP36L1 interactions, we detected interactions between ZFP36L1 and the phosphoserine-containing protein binders YWHAB/Z (**Figure 3A**). These 14-3-3 proteins were previously described to regulate the protein turnover of ZFP36L1 ^66,67^. In addition, ZFP36L1 interacted with the protein quality control protein SQSTM1 (p62), which relocates ubiquitinylated proteins to the lysosome for degradation (**Figure 3A**). Lastly, ZFP36L1 interacts with SIRT2 (**Figure 3A**), which de-acetylates lysines and thereby alters protein expression and function ^68^. In conclusion, UltraID proximity labeling uncovered a multitude of ZFP36L1 interactions with proteins that contribute to a broad range of regulatory pathways, each defining different aspects of regulating the fate of target mRNA and protein expression.

### Time-dependent protein-protein interactions with ZFP36L1

Having identified a broad scala of proteins in close proximity with ZFP36L1 (**Figure 3**), we next questioned whether these interactions were constant, or whether they were subject to change over the course of T cell activation. To answer this question, we performed a time-lapse proximity labeling, by activating T cells for 2h, 5h, and 16h, and by adding biotin between these timepoints for 2h, for 3h (from 2-5h of activation) or for 9h (from 5-16h of activation) (**Figure 4A, Supplementary Figure 4A)**. These time points were chosen based on the effects on previously measured target mRNAs of ZFP36L1, which can occur early on (2h), at 5h of T cell activation, or rather later at 16h of T cell activation (Popovic 2023, Petkau 2022, Zandhuis 2025).

**Figure 4.**
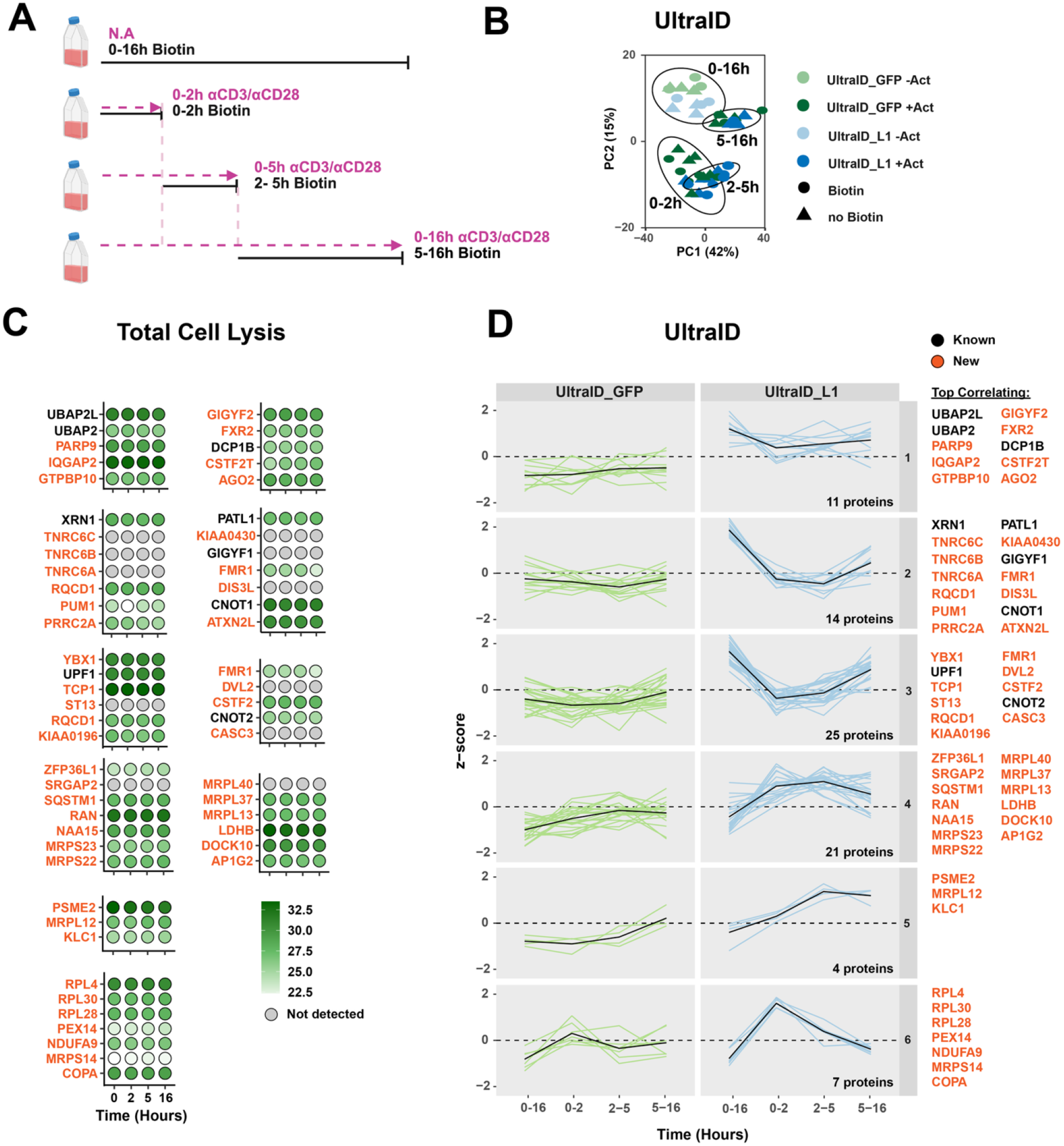
Time-resolved proximity labeling reveals dynamic interactions with ZFP36L1 during T cell activation. **a**. Graphical representation of UltraID time course experiment. T cells were re-activated with aCD3/aCD28 for indicated time points (magenta dotted arrow) in the presence or absence of biotin. Biotin was added at indicated time points (black line). **b**. PCA for indicated conditions. Outlined circles indicate different timepoints of T cell activation. Color indicates UltraID_L1 (Blue) and UltraID_GFP (Green). Circle indicates the addition of biotin, triangle: no biotin added. **c**. Protein abundance in full cell lysates of proteins that were significantly enriched for biotinylation (LFC>1, p.adj <0.05) in the time course of UltraID_L1 samples (panel d). Grey indicates not detected. Fold-change (log2FC) enrichment is indicated by color code. n= 3 donor pools. **d**. Scaled intensity profiles of UltraID_L1-enriched proteins across activation time points, as determined by supervised classification of 6 shapes (Supplementary Figure 4B) identified to contain interaction partners with a Pearson correlation of >0.5, and p.adj <0.05 and LFC>1. n= 3 donor pools. *See Supplementary Figure 4*

PCA of biotinylated proteins revealed that the time of T cell activation rather than the biotinylation time was the prime driver of differential protein abundances, and in addition that the UltraID_L1 pulldowns slightly clustered away from UltraID_GFP control pulldowns (**Figure 4B; Supplementary Table 8**). Supervised classification of binding kinetics of proteins in proximity to ZFP36L1 (LFC>1, adjusted p<0.05, Pearson correlation ≥0.5) resulted in 11 shapes, of which 6 shapes contained enriched proteins (**Supplementary Figure 4B, C; Supplementary Table 8, 9)**. MS analysis of total cell lysates revealed limited changes throughout T cell activation of protein abundance of ZFP36L1 interactors, thereby excluding the possibility that alterations of interactions are a mere consequence of differential protein levels **(Figure 4C, Supplementary Table 10)**.

Many proteins in proximity to ZFP36L1 were already detected in resting T cells (0h; **Supplementary Figure 4C)**, which points to robust, activation-independent interactions with ZFP36L1. Alternatively, some interactions may also result from the exogenous ZFP36L1 expression in UltraID_L1-containing T cells, which is slightly higher than that of endogenous ZFP36L1 (**Figure 1D**). Nevertheless, we could clearly identify differential protein interaction profiles with ZFP36L1 throughout the 16h time course of T cell activation. Some proteins such as the SG/PB-associated proteins UBAP2 and UBAP2L, the decapping protein DCP1B, as well as GIGYF2 and CSTF2T interacted with ZFP36L1 throughout T cell activation (shape 1; **Figure 4D**). Other proteins reduced their interaction with ZFP36L1 at 2h before they gradually increased their interactions again (shapes 2+3, **Figure 4D**). This class included proteins involved in mRNA degradation, and translation repression, such as including UPF1, GIGYF1, and CSTF2, but also the deadenylation proteins TNRC6A-C and the SG/PB-associated proteins ATXN2L, PUM1, PATL1, and YBX1 **(Figure 4D)**.

We also identified proteins that increased their interaction with ZFP36L1 upon T cell activation (shapes 4+5; **Figure 4D)**. Intriguingly, the majority of these were mitochondrial small and large ribosomal proteins that are translated in the cytoplasm, after which they are transported to the mitochondria where they assemble into complexes^69^ (MRPS & MRPL, **Figure 4D**). T cell activation triggers mitochondrial biogenesis, which is essential for translating cytotoxic molecules and for maintaining sustained cytotoxicity ^70–72^. Because the total protein levels of mitochondrial proteins did not change upon T cell activation **(Figure 4C, Supplementary Table 10)**, ZFP36L1 may thus potentially regulate their localization and/or transport of translated proteins.

We also identified proteins that only briefly increased their interaction with ZFP36L1 at 2h of activation (shape 6, **Figure 4D**). These included the ribosomal proteins RPL4, RPL28 and RPL30, and the mitochondrial proteins MRPS14 and NDUFA9, the latter being a component of Complex I of the respiratory chain ^73^ (**Figure 4D**). In conclusion, ZFP36L1 displays both constant and time-dependent interactions, and these altered interactions belong to different RNA regulatory pathways, which could define different emphasis of ZFP36L1 on its divergent PTR functions over the course of T cell activation.

### RNA-dependent and -independent interactions with ZFP36L1

Having deciphered the ZFP36L1 interactome and its kinetics, we next asked whether the interactions with ZFP36L1 required the presence of RNA. To study this, we performed a co-immuno-precipitation (Co-IP) of ZFP36L1 with untreated T cell lysates or with RNAse pre-treated lysates (**Supplementary Figure 5A)**. MS analysis revealed that in addition to ZFP36L1 enrichment versus IgG control, treatment with RNAse defined the distribution of samples in the PCA **(Figure 5A, Supplementary 5B, C; Supplementary Table 11, 12**; LFC≥1; p.adj<0.05).

**Figure 5.**
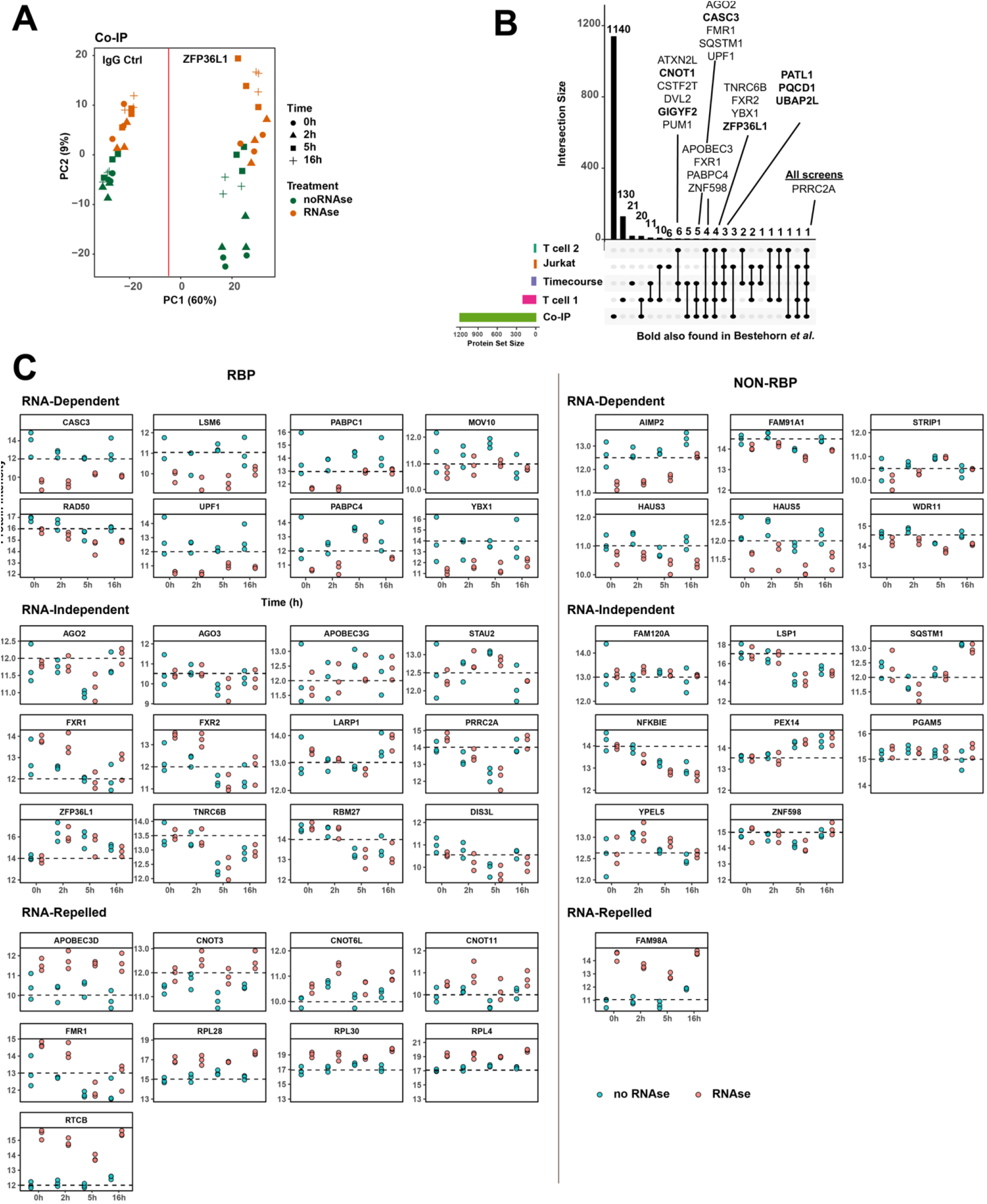
RNA-dependency of interactions with ZFP36L1. Co-IP with ZFP36L1 antibodies and IgG control with T cells lysates generated at indicated time points that were treated or not with 200 U/mL RNAse A during the entire protocol (lysis, incubation & washes). a. PCA of indicated conditions. Green dots: RNA present, Orange dots: RNAse A treated. Different shapes represent different timepoints of activation. **b**. Upset plot of proteins identified as ZFP36L1 interactors in the Co-IP or indicated proximity labeling essay. Numbers refer to numbers of proteins found interacting with ZFP36L1 in indicated conditions. LFC>1, p.adj<0.05, shape correlation >0.5. Proteins indicated in bold were also identified in HEK293T cells^42^. **c**. LFQ protein intensities from Co-IP time course of 44 proteins co-identified by the Co-IP and at least 1 proximity labeling (panel b). Green dots: RNA present, Orange dots: RNAse A treated. Proteins were categorized into known RNA-binding proteins (RBPs) and non-RBPs that displayed RNA-dependent, RNA-independent or RNA-repelled interactions with ZFP36L1 (as defined in Supplementary Figure 5F). n= 3 donor pools. *See Supplementary Figure 5*

The Co-IP executed in the presence of RNA identified several prototypic ZFP36L1 interactors, including the CCR4-NOT and EXOCS proteins, together with its sister proteins ZFP36 and ZFP36L2 (**Supplementary Figure 5C)**. In line with the proximity labeling time course **(Figure 4)**, the Co-IP also detected ZFP36L1 interactome dynamics throughout T cell activation, with most differential interactions measured at 5h of T cell activation **(Supplementary Figure 5D)**. The majority of protein interactions with ZPF36L1 did not change with pre-treatment and were thus RNA independent, as exemplified for 0h and 5h of T cell activation **(Supplementary Figure 5E**). Yet, some interactions were reduced upon RNAse treatment (RNA-dependent) or even increased (RNA-repelled; **Supplementary Figure 5E**). Because the overlap of ZFP36L1 interaction partners was limited between the Co-IP and proximity labeling experiments **(Figure 5B)**, we focused for downstream analysis on the 44 proteins that were co-identified in at least 1 UltraID screen, and that we therefore considered high-confidence ZFP36L1 interactors **(Supplementary Figure 5F)**. RBPs that displayed reduced interactions with ZFP36L1 in the absence of RNA included UPF1 ^50,51^, YBX1, MOV10, the exon junction complex protein CASC3 ^74^, LSM6, a component of the RNA degradation and spliceosome complex ^75,76^, and the poly-A binding proteins PABPC1 and PABPC4 ^77,78^ **(Figure 5C: *RNA-dependent*; Supplementary Table 11)**. The preferred interaction with ZFP36L1 in the presence of RNA was constant throughout time for all proteins, except for MOV10, which showed the highest RNA-mediated ZFP36L1 interaction at 5h of T cell activation **(Figure 5C)**.

Non-RBPs preferentially interacting with ZFP36L1 in the presence of RNA were AIMP2, **a** scaffold for the tRNA synthetase complex^79,80^, in addition to WDR11 and FAM91A1, which belong to the complex that tethers vesicles to the Golgi apparatus ^81^. We also identified HAUS3 and HAUS5 involved in microtubule generation^82,83^ and STRIP1 (Fam40a), which regulates the cell shape by modulating actin filament dynamics ^84,85^, the latter with strongest effects of RNA depletion at 0h and 2h of T cell activation **(Figure 5C – RNA-dependent)**. Interestingly, CNOT proteins showed increased binding to ZFP36L1 when RNA was depleted **(Figure 5C RNA-repelled)**, suggesting that the interaction to the c-terminal CNOT interacting motif of ZFP36 proteins ^36,38^ does not require the presence of RNA to maintain its interaction with ZFP36L1.

Some RBPs were oblivious to RNA depletion for their interaction with ZFP36L1 (AGO2, AGO3, APOBEC3G, Stau2, FXR1, FXR2, LARP1, PRRC2A, TNCRB6B, RBM27, DISL3 and ZFP36L1) **(Figure 5C; RNA-independent)**. As expected, interactions with SQSTM1 (p62) and proteins involved in protein turnover and post-translational modifications did not require RNA (FAM120A, LSP1, NFKBIE, PEX14, PGAM5, YPEL5, the scaffolding protein ZNF598), **Figure 5C**). Fam98A, which stimulates PRMT1-induced arginine methylation showed even increased interaction with ZFP36L1 when RNA was depleted ^86^ **(Figure 5C**). In conclusion, this analysis uncovers the context-dependent nature of interactions with ZFP36L1.

### UPF1 KO and GIGYF1/2 KO T cells display cellular stress through increased protein translation

Lastly, we aimed to decipher the role of ZFP36L1 candidate interactors in T cells. We focused on 4 proteins that exert divergent post-transcriptional functions: the nonsense-mediated decay factor UPF1, the de-adenylation proteins/translational repressors GIGYF1 and GIGYF2, and the nuclear alternative poly-adenylation regulating protein CSTF2T. UPF1 interacts with ZFP36L1 preferentially at 0h and 16h, and it does so in an RNA-dependent manner (**Figure 4D/5C**). GIGYF1 shows similar interaction kinetics with ZFP36L1 as UPF1 (**Figure 4D**). Conversely, GIGYF2 and CSTF2T maintain their interaction with ZFP36L1 throughout T cell activation (**Figure 4D**). Of note, UPF1, GIGYF1 and GIGYF2 belong to the core ZFP36L1 interactome, being identified also in HEK293T cells^42^ (**Supplementary Figure 2, 5 & 7**).

We first gene-edited T cells with CRISPR-Cas9 to delete UPF1, CSTF2T, GIGYF1 and GIGYF2 (**Supplementary Figure 6A)**. Because GIGYF1 and GIGYF2 cooperate with each other ^87^ and single deletions had no effect (not shown), we generated GIGYF1/GIGYF2 double deficient T cells (GIGYF1/2 KO). The viability of GIGYF1/2 KO and CSTF2T KO T cells was comparable to that of control T cells (**Supplementary Figure 6B)**. Only UPF1 KO T cells were less viable **(Supplementary Figure 6B)**. We next measured the global translation rate with puromycin incorporation for 10 min. CSTF2T deficiency showed very limited effects in resting and in activated T cells (**Figure 6A)**. In contrast, UPF1 KO and GIGFYF1/2 KO T cells displayed substantially higher global translation rates in resting T cells compared to control T cells (**Figure 6A**). T cell activation for 2h led to a substantial drop in puromycin incorporation in both UPF1 KO and to some extent GIGFYF1/2 KO T cells, to levels that were lower than those of control T cells (**Figure 6)**. This could not merely be attributed to cell viability for GIGYF1/2 KO T cells **(Supplementary Figure 6B);** although UPF1 KO displayed high cell death upon activation. Because ZFP36L1 KO T cells showed very limited changes in puromycin incorporation **(Supplementary Figure 6C)**, we conclude that UPF1 and GIGYF1/2 are the driving force in changing the global translation rates.

**Figure 6.**
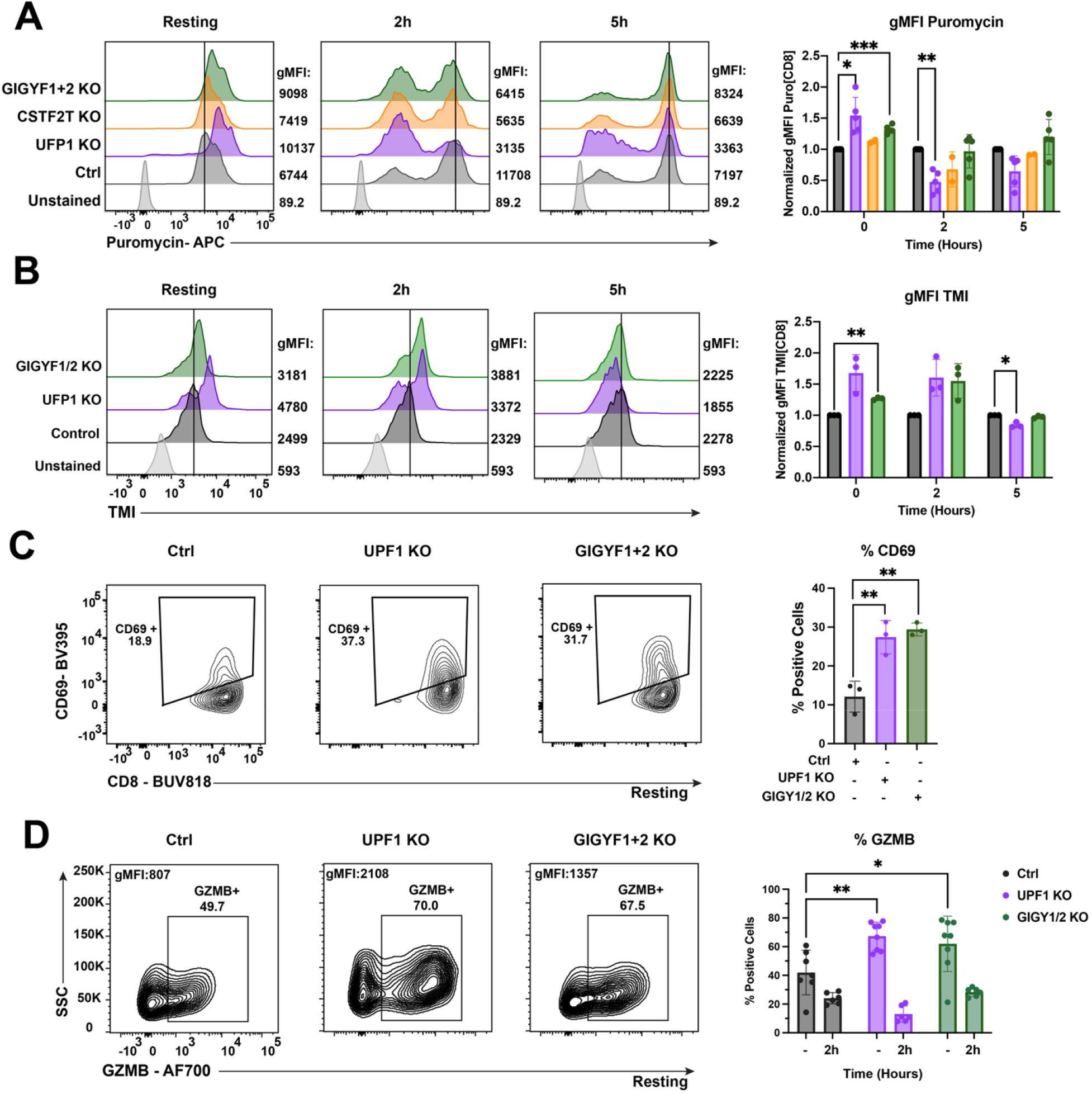
UPF1 and GIGYF1/2 deletion alters translation and stress responses in human CD8^+^ T cells. **a**. Puromycin incorporation in indicated CRISPR-Cas9 gene-edited T cells or non-targeting crRNA (ctrl)-edited T cells at indicated time points of activation with aCD3/aCD28. Left panel: Representative plot of puromycin incorporation as quantified by flow cytometry. Geometric Mean Fluorescent Intensity (gMFI) value indicated on the right. Right panel: puromycin gMFI during rest or indicated time of stimulation with aCD3/aCD28. n=2-5 donors. **b**. Detection of unfolded proteins with tetraphenyl ethene maleimide (TMI) dye. Left panel: Representative plot of TMI as quantified by flow cytometry. Geometric Mean Fluorescent Intensity (gMFI) value indicated on the right. Right panel: TMI gMFI during rest or indicated time of stimulation with aCD3/aCD28. n= 3 donors. **c**. Left panel: representative CD69 expression in resting control and indicated KO T cells measured by flow cytometry. Right panel: percentage of CD69^+^ T cells. Data shown as mean ± SD. n=3 donors. **d**. Left panel: Representative Granzyme B (GZMB) expression measured by flow cytometry. Right panel: percentage of GZMB + T cells in resting conditions and after 2h of stimulation with aCD3/aCD28. n=6 donors, data shown as mean ± SD (right). A& B) Two-way ANOVA with Dunnett’s correction C& D) One-way ANOVA with Dunnett’s correction. *p < 0.05, ** p < 0.01, *** p<0.001. *See Supplementary Figure 6*

Higher protein production can lead to unfolded protein response and consequentially, induce cell stress^88^. We therefore measured the level of unfolded proteins with tetraphenyl ethene maleimide (TMI), which becomes fluorescent when engaging with free thiol side chains that are exposed in unfolded proteins ^89^. Concomitant with lower cell viability, UPF1 KO T cells showed increased levels of unfolded proteins (**Figure 6B**), Also GIGYF1/2 KO cells had higher levels of TMI staining at resting state, and particularly at 2h of T cell activation, although not statistically significant **(Figure 6B)**. After 5h of activation, however, the levels of unfolded proteins normalized for all control and KO T cells (**Figure 6B**).

T cell activation substantially increases the translation rate (Wolf et al. 2020; Marchingo et al. 2020; Sinclair and Cantrell). We therefore questioned whether the observed higher puromycin incorporation in resting UPF1 KO and GIGYF1/2 KO T cells indicated an altered T cell activation status. Indeed, the percentage of resting T cells expressing the activation marker CD69 more than doubled in UPF1 KO and GIGYF1/2 KO T cells compared to control T cells **(Figure 6C)**. Granzyme B expression was also substantially higher in resting UPF1 KO and GIGYF1/2 KO T cells, and T cell activation resulted in stronger release of Granzyme B (=loss of signal), most prominently in UPF1 KO T cells, again pointing to a high T cell activation state in the absence of T cell triggers (**Figure 6D)**. Combined, these data suggest that UPF1 and GIGYF1/2 contribute to T cell fitness by limiting protein translation and activation status in resting T cells.

### UPF1 promotes ZFP36L1 protein production

We next sought to decipher the interplay of ZFP36L1 with UPF1 and GIGYF1/2. Re-analysis of the comparative interactome study of wild type ZFP36L1 with a tandem zinc finger (TZF) mutant failing to interact with RNA by Bestehorn et al. ^42^ revealed that only GIGYF1, but not UPF1 and GIGYF2, lost its interaction with ZFP36L1 when the TZF was mutated (**Figure 7A**), indicating divergent requirements of these three proteins for interacting with ZFP36L1.

**Figure 7.**
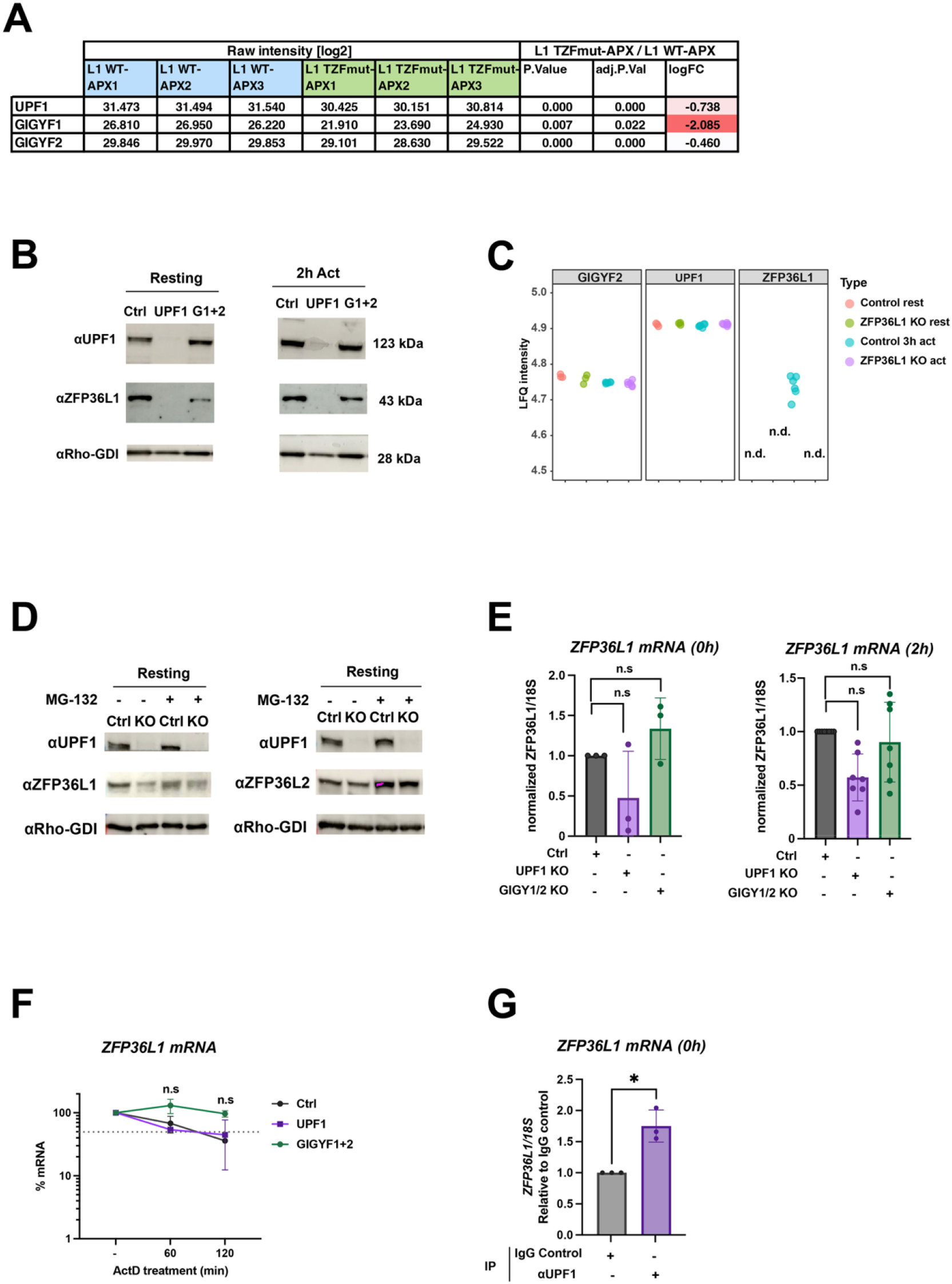
UPF1 interacts with ZFP36L1 mRNA and modulates its expression at the transcript level. **a**. Quantification of ZFP36L1 interactors identified by immunoprecipitation–mass spectrometry (IP-MS) in HEK293T cells expressing either wild-type or zinc-finger mutant (TZFmut) ZFP36L1. Data extracted from Bestehorn *et al*.^42^ **b**. Immunoblot of ZFP36L1 and UPF1 in non-targeting crRNA (Ctrl) treated, UPF1 KO (UPF1) and GIGYF1+2 KO (G1+2) from resting or aCD3/aCD28 activated T cells (2h). aRho-GDI was used as loading control. Representative of at least 6 donors. **c**. WT and ZFP36L1 KO cells were analysed using MS. LFQ intensity of GIGYF2, UPF1 and ZFP36L1 from human control and ZFP36L1 gene edited T cells, extracted from MS data depicted at either resting state or 3h PMA/Iono activations. n= 6 independent donors. n.d = not detected. **d**. Immunoblot of ZFP36L1, ZFP36L2 and UPF1 in control and UPF1 KO (KO) T cells treated with MG132 or vehicle control for 2h, executed in resting T cells (left) or 2h aCD3/aCD28 re-activated T cells (right). **e**. *ZFP36L1* mRNA levels in control, UPF1 KO and GIGYF1+2 KO cells at resting state (left) and 2h of activation (right), measured by RT-qPCR. Data normalized to *18S*. **f**. RNA stability assay of *ZFP36L1* using actinomycin D treatment started after 2h of activation. mRNA decay measurements of *ZFP36L1* mRNA. **g**. Native RNA immunoprecipitation (RIP) with UPF1 antibody or IgG isotype control from lysates of resting T cells. qRT-PCR of endogenous mRNAs from RIP of UPF1 relative to IgG control. n=3 donor pools, mean ± SD. E-G) Two-way ANOVA with Dunnett’s correction. *p < 0.05. n.s = non-significant.

Strikingly, when we measured the effect of UPF1 and GIGYF1/2 depletion on ZFP36L1 protein expression, we found that UPF1 deletion substantially reduced the ZFP36L1 protein levels in both resting and activated T cells (2h; **Figure 7B)**. Because neither UPF1 nor GIGYF2 protein expression was altered in ZFP36L1 KO T cells **(Figure 7C)**, we conclude that UPF1 is an upstream regulator of ZFP36L1 protein expression. UPF1 does not modulate the protein stability of ZFP36L1, because blocking the proteasomal degradation with MG132 did not recover ZFP36L1 protein expression in UPF1 KO T cells (**Figure 7D**). In contrast, proteasomal block restored the protein expression of its sister protein ZFP36L2, irrespective of UPF1 deletion (**Figure 7D)**. Thus, we concluded that UPF1 regulates the generation of ZFP36L1 protein.

Protein output is in part defined by the number of mRNA templates. Interestingly, ZFP36L1 mRNA expression was reduced in UPF1 KO T cells, but not in GIGYF1/2 KO T cells, yet without overt effects on mRNA stability during the 2h treatment with the transcription inhibitor Actinomycin D **(Figure 7E, F)**. A native RNA-immunoprecipitation followed by qRT-PCR (RIP-qRT-PCR) on unmodified T cells confirmed that UPF1 interacts with *ZFP36L1* mRNA **(Figure 7G)**, further supporting the hypothesis that UPF1 is an upstream regulator of ZFP36L1 expression. Thus, UPF1 not only interacts with ZFP36L1 protein as uncovered by proximity labeling, but UPF1 also supports the protein expression of ZFP36L1, at least in part by regulating its transcript levels, and we hypothesize that the helicase activity of UPF1 could potentially contribute to this. Combined, this finding showcases how proximity labeling can uncover novel protein-protein interactions and help identify novel regulatory nodes of proteins, as exemplified here for ZFP36L1.

## DISCUSSION

In this study, we uncovered the multifaceted nature of the ZFP36L1 interactome in human T cells. We report versatile interaction kinetics with ZFP36L1 during T cell activation, and we defined how RNA mediates these interactions. This comprehensive interactome analysis positions ZFP36L1 at the center of multiple post-transcriptional regulatory pathways in T cells.

Beyond its established role in recruiting the CCR4-NOT deadenylation complex ^36–39^, we detected proximity of ZFP36L1 to a multitude of novel interactions with proteins involved in mRNA decapping (PATL1 and DDX6) and mRNA degradation pathways, as well as translation repressors (GIGYF1, GIGYF2, ZNF598 and 4EHP) and other regulatory pathways. Notably, many of the newly identified candidate interactors are part of complexes of previously annotated ZFP36L1 interactors. Therefore, it would be interesting to decipher how ZFP36L1 comes in proximity with the different subunits, and to determine whether ZFP36L1 plays an active role in the formation of these RBP complexes. One way we aimed to decipher this was by defining the requirement of RNA for interacting with ZFP36L1. These data clearly depicted the RNA (in)dependency of several candidates. However, it should also be considered that some protein-protein interactions indicated as RNA-independent may initially be mediated by RNA and persist due to high affinity interactions or conformational locking.

A key finding of our study is that the ZFP36L1 interactome undergoes remodeling during T cell activation. For instance, proteins involved in mRNA degradation and translation repression processes are reduced during early T cell activation. We hypothesize that this temporary suspension of these interactions may be required to allow for efficient protein synthesis to occur, a key feature to drive rapid remodeling of the T cell proteome upon T cell activation ^1–3^. The subsequent re-establishment of these interactions at 5h of T cell activation coincides with the time point when ZFP36L1 commences to dampen the T cell activation program, including for instance the cytokine production ^17,23,24^. How the landscape of interaction partners is altered remains to be established. Because we find very limited alterations in RBP expression levels upon T cell activation, it is conceivable that post-translational modifications (PTMs) define the alterations in the RBP interactome of ZFP36L1 to allow for the rapid shift in the proteome and production of effector molecules. Supporting this hypothesis, we identified several post-translational modifiers and PTM-binders interacting with ZFP36L1, including the 14-3-3 proteins YWHAB/YWHAZ interacting with post-translational modifications, the protein deadenlyase SIRT2, and the CTLH E3 ubiquitin protein ligase complex component YELP5 (**Figure 3**). Future studies should address which signaling pathways downstream of the T cell receptor and costimulatory molecules drive these modifications and thus alter ZFP36L1’s interactions and its activity. In conclusion, the time-lapse proximity labeling during T cell activation demonstrates that the ZFP36L1 interactome is context dependent and versatile.

The advantage of proximity labeling is that it can detect interactions in living cells, capturing physiologically relevant interactions in their native cellular context, including weak and transient interactions^91,92^. In particular transient interactions are typically lost during cell lysis procedures required for e.g. Co-IP. For instance, whereas GIGYF1, GIGYF2, and CSTF2T interactions were clearly measured with proximity labeling, we failed to detect these interactions over background in the ZFP36L1 Co-IP. Furthermore, proximity labeling identified interactions of ZFP36L1 with nuclear factors such as the polyadenylation factors CSTF2 and CSTF2T, and the RNA decapping protein PATL1, which can shuttle between nucleus and cytoplasm ^93^. Moreover, we also identified proteins in proximity to ZFP36L1 that are associated with granules (G3BP1 FXR1/2, ATXN2L, PUM1, YBX1, UBAP2/L, MOV10)^61^, structures, or membrane-bound organelles (NUPL1; nuclear pore complex ^94^), and these cellular compartments are generally disrupted during cell lysis. This included also interactions with non-RBPs, like SIRT2, YELP5 and SQSTM1.

Our investigation of functional interactions also revealed a previously uncharacterized relationship between UPF1 and ZFP36L1. UPF1, a central component of the nonsense-mediated mRNA decay pathway ^50,51,95,96^, interacts with ZFP36L1 protein primarily in resting T cells and at later time points of T cell activation when translation activity is returning back towards levels of resting T cells. UPF1 orchestrates co-translational mRNA surveillance by recruiting the endonuclease SMG6 and UPF2 ^97^. These two proteins were not identified by proximity labeling as ZFP36L1 interactors with SMG6 or UPF2. Therefore, it remains to be determined whether the helicase activity of UPF1 supports the entry of ZFP36L1 to its target mRNA together with SMG6 and UPF2, or whether it exerts a fundamentally different activity with ZFP36L1.

UPF1, however, also appears to directly regulate the protein expression of ZFP36L1. ZFP36L1 protein expression is rapidly induced upon T cell activation ^17,23,24^. In the absence of UPF1, ZFP36L1 protein expression in resting and activated T cells is substantially impaired. UPF1 binds to ZFP36L1 mRNA, and its depletion reduces the overall mRNA levels of ZFP36L1 mRNA. These findings combined identify UPF1 as an upstream regulator of ZFP36L1 protein expression, and they suggest that UPF1 possibly monitors and thereby safeguards the quality of ZFP36L1 transcripts, allowing for its protein production to occur. This, in combination with the evidence for ZFP36L1:UPF1 interaction, could implicate that ZFP36L1 can indirectly regulate its own levels through UPF1.

In conclusion, our findings position ZFP36L1 as a central coordinator in a complex network of post-transcriptional regulatory mechanisms in T cells. Through its dynamic interactions with components of mRNA decay, translation repression, and potentially mitochondrial translation machineries, ZFP36L1 appears equipped to orchestrate the temporal sequence of gene/protein expression events required for proper T cell activation. The regulatory relationship between UPF1 and ZFP36L1 highlights the intricate interdependence of post-transcriptional control systems and creates new opportunities for understanding how these mechanisms shape T cell function in immunity and disease.

## MATERIALS AND METHODS

### Cell culture

Peripheral blood mononuclear cells (PBMCs) from anonymized healthy donors were used in accordance with the Declaration of Helsinki (Seventh Revision, 2013) after written informed consent (Sanquin). PBMCs were isolated by Lymphoprep density gradient separation (Stemcell Technologies) and cryopreserved until further use. Human T cells and Jurkat cells were cultured in Culture Medium (Iscove’s Modified Dubecco’s Medium (IMDM) supplemented with 10% fetal bovine serum (FBS), 100 U/mL penicillin, 100 µg/mL streptomycin, and 2 mM L-glutamine supplemented with 100 IU/mL recombinant human rhIL2 (Protech) and 10 ng/mL rhIL-15 (Protech). Of note, because biotin deprivation induces stress and inflammatory response in T cells (Elahi, Sabui, and Said 2018), T cells were cultured in biotin-containing media (53nM). Human fibrosarcoma FLYRD18 cells (ADCC) were cultured in IMDM supplemented with 10% FBS, 1% L-glutamine, 1% Penicillin-Streptomycin antibiotic solution at 37°C, 5% CO2 and split every 2 days at 1:10. All cells were cultured at 37°C with 5% CO_2_.

### T cell activation

T cells from defrosted PBMCs were activated for 48h as previously described ^98^. Briefly, 24-well plates were pre-coated overnight at 4°C with 4µg/mL rat a-mouse IgG2a (MW1483, Sanquin) in phosphate-buffered saline (PBS). Plates were washed with PBS and coated for >3 h with 1 µg/mL αCD3 (HIT3a, Biolegend) at 37°C. 1.3×10^6^ PBMCs/well were seeded with 1 µg/mL soluble αCD28 (CD28.2, Biolegend) in 1 mL Iscove’s Modified Dubecco’s Medium (IMDM) supplemented with 10% fetal bovine serum (FBS), 100 U/mL penicillin, 100 µg/mL streptomycin, and 2 mM L-glutamine. After 48h of activation at 37°C, 5% CO2, cells were harvested and cultured in standing T25/T75 tissue culture flasks (Thermo Scientific) at a density of 0.8×10^6^/mL in culture medium. Media was refreshed every 2-3 days. Upon nucleofection, T cells were cultured in T cell mixed media (Miltenyi) supplemented with 5% heat-inactivated human serum, 5% FBS, 100 U/mL Penicillin, 100 µg/mL streptomycin, 2 mM L-glutamine, 100 IU/mL rhIL-2 (Peprotech), and 10 ng/mL rhIL-15 (Peprotech) for 4 days after switching back to Culture Medium for another 3 days. T cells were re-activated in Culture Medium in 96-wells plates with 10 ng/mL Phorbol Myristate Acetate (PMA) and 1μM Ionomycin, or with αCD3 (Pelicluster; Sanquin) and αCD28 (Biolegend), 1µg/mL each. For proteasomal inhibition, 2µM of MG-132 (Sigma-Aldrich) was used.

### Generation UltraID constructs

UltraID_ZFP36L1_GFP or UltraID_GFP fusion proteins were generated as gBlock DNA by IDT. These were then transferred into the pMX_Puro backbone (Addgene) into ClaI and EcoRI restriction sites according to the manufacturer’s protocol (T4; NEB) (sequence upon request). pMX_UltraID was generated by digesting pMX_UltarID_ZFP36L1_P2A_GFP vector with NotI and ligating it back together to remove ZFP36L1_P2A_GFP. Plasmid DNA was amplified in DH5*α* super-competent bacteria (NEB) and isolated with NucleoSpin Plasmid Transfection-grade Mini-Prep isolation kit (Machery-Nagel). Sequences were confirmed with Sanger sequencing.

### Virus production and retroviral transduction

1×10^5^ FLYRD18 cells/well were pre-seeded in 6-well culture plates and cultured overnight at 37°C. Transfection was carried out with GeneJammer (Agilent) according to the manufacturer’s protocol. Cells were cultured for 48h at 32°C. αCD3/αCD28-activated **T** cells (48h) were retrovirally transduced as previously described^24^. Briefly, non-tissue culture treated 24-well plates were pre-coated overnight with 50 µg/mL Retronectin (Takara). Plates were washed once with PBS, 800 µL viral supernatant/well was added and spun at 4°C at 2820g. Supernatant was removed and 1×10^6^ T cells/well were added, plates spun for 5 min at 180g and incubated overnight at 37°C. Supernatant was replaced with fresh Culture Medium. After 48h, T cells were harvested and cultured in T25/T75 flasks for 6–8 days as described above.

### Flow cytometry

To analyze GFP protein expression, T cells were washed with FACS buffer (PBS, containing 1% FBS and 2 mM EDTA) and labeled for 20 min at 4°C with indicated antibodies (see Appendix A). Dead cells were excluded with Near-IR (Life Technologies). For intracellular cytokine staining, cells were cultured with 1 µg/mL brefeldin A for indicated time points, fixed, and permeabilized with Cytofix/Cytoperm kit according to the manufacturer’s protocol (BD Biosciences) prior to acquisition using FACSymphony. Data were analyzed with FlowJo (BD Biosciences, version 10).

### Immunoblotting

Cell lysates (1×10^6^ cells) were prepared by standard procedures using RIPA lysis buffer (Thermo) supplemented with protease and phosphatase inhibitors (Thermo). Lysates were run on 4–12% SDS-PAGE (Thermo). SDS-PAGE gels were directly transferred onto nitrocellulose membranes (iBlot2, Thermo). Membranes were blocked with 5% BSA TBST solution (Fraction V, Sigma) and incubated overnight with α-Streptavidin-HRP, α-UPF1 (Proteintech), α-CSTF2T (Proteintech), ZFP36L1 (abcam) or α-RhoGDI (MAB9959, Abnova), followed by α-Rabbit (4050-05, Southern Biotech) or α-Mouse (1031-05, Southern Biotech) HRP-conjugated secondary antibodies.

### UltraID

Proximity-dependent biotin identification assay was performed according to Kubitz et al.^30^, with the following modifications. Retrovirally transduced Jurkat cells and human T cells were FACS-sorted based on GFP expression 3 days post transduction and expanded for another 7 days. Cells were activated with αCD3/αCD28 for 2-16 h or left unactivated (rested) in the presence or absence of (50-) 500µM biotin dissolved in DMSO and incubated in the dark at 37°C. Cells were harvested, washed twice with ice-cold PBS, and lysed in 1 ml lysis buffer (10 mM Tris/HCl pH 7.5, 150 mM NaCl, 0.5 mM EDTA + HALT Protease/Phosphatase Inhibitor cocktail (Thermo, 1861281) + 0.5% NP40 (v/v)) for 5 min in at 4 °C. Lysates were sonicated 3 x 30 sec using a Branson Sonifier 450 device, keeping cells on ice for 1 min in between sessions. Cell lysates were cleared by centrifugation 14,000xg at 4°C for 20 min. Cleared lysates were added to pre-washed 200 μl Dynabeads MyOne Streptavidin C1 (65002, Invitrogen) beads (2x wash with PBS, equilibrated in lysis buffer for 5 min at 4°C and washed with lysis buffer). Mix was incubated overnight rotating at 4°C. The following day, beads were washed 3x using wash buffer 1 (10 mM Tris/HCl pH 7.5, 150 mM NaCl, 0.5 mM EDTA + HALT Protease/Phosphatase Inhibitor cocktail (Thermo, 1861281) and wash buffer 2 (10 mM Tris/HCl pH 7.5, 150 mM NaCl). Proteins were digested on-bead and prepared for Mass Spectrometry analysis as described below.

### Co-immunoprecipitation

Cytoplasmic lysates of 50×10^6^ αCD3/αCD28-activated human CD3+ T cells were prepared (simultaneously prepared from same donor pools cells of the UltraID timecourse experiment of **Figure 4)** using lysis buffer (140 mM NaCl, 5 mM MgCl2, 20 mM Tris/HCl pH7.6, 1% Digitonin) freshly supplemented with 1% protease inhibitor cocktail (Sigma). Protein A Dynabeads (Invitrogen) were prepared according to the manufacturer’s protocol. Cell lysate was pretreated with 200 U/mL RNAse A (Thermo Scientific). immunoprecipitated for 4h at 4ºC with 10 µg polyclonal rabbit a-ZFP36L1 (ABN192, Sigma-Aldrich) or with isotype control (12–370, Sigma-Aldrich) ± 200 U/mL RNAse A (Thermo Scientific). Beads were washed twice with wash buffer (150 mM NaCl, 10 mM Tris/HCl pH7.6, 2 mM EDTA, protease/phosphatase inhibitor cocktail) and twice with 10 mM Tris/HCl pH7.6. Immunoprecipitated proteins were reduced and on-bead alkylated. Proteins were detached with 250 ng trypsin for 2 h at 20ºC. Beads were removed and proteins were further digested into peptides with 350 ng trypsin for 16 h. Peptides were prepared for MS analysis, as described below.

### Mass spectrometry data acquisition

For the UltraID T cell screen 1 and Jurkat screen: tryptic peptides were separated by nanoscale C18 reverse phase chromatography coupled online to an Orbitrap Fusion Lumos Tribrid mass spectrometer (Thermo Scientific) via a nanoelectrospray ion source (Nanospray Flex Ion Source, Thermo Scientific). Peptides were loaded on a 20cm 75–360µm inner-outer diameter fused silica emitter (New Objective) packed in-house with ReproSil-Pur C18-AQ, 1.9μm resin (Dr Maisch GmbH). The column was installed on a Dionex Ultimate3000 RSLC nanoSystem (Thermo Scientific) using a MicroTee union formatted for 360μm outer diameter columns (IDEX) and a liquid junction. The spray voltage was set to 2.15kV. Buffer A was composed of 0.1% formic acid and buffer B of 0.1% formic acid, 80% acetonitrile. Peptides were loaded for 17 min at 300nl/min at 5% buffer B, equilibrated for 5 min at 5% buffer B (17-22 min) and eluted by increasing buffer B from 5-28% (22-80 min) and 28-40%(80-85 min), followed by a 5 min wash to 95 % and a 5 min regeneration to 5%. Survey scans of peptide precursors from 375 to 1500 m/z were performed at 120K resolution (at 200 m/z) with a 4×10^5^ ion count target. Tandem mass spectrometry was performed by isolation with the quadrupole with isolation window 0.7, HCD fragmentation with normalized collision energy of 30, and rapid scan mass spectrometry analysis in the ion trap. The MS2 ion count target was set to 3×10^4^, and the max injection time was 20ms. Only those precursors with charge state 2–7 were sampled for MS2. The dynamic exclusion duration was set to 20s with a 10ppm tolerance around the selected precursor and its isotopes. Monoisotopic precursor selection was turned on. The instrument was run in top speed mode with 1s cycles. All data were acquired with Xcalibur SII software^99^. For the UltraID T cell sreen 2, UltraID timecourse screen, and ZFP36L1 Co-IP, samples were measured using TimsTOF-DIA as previously described ^100^. One amendment was made to the protocol, which encompassed that 30 samples per day were measured instead of 60. For the total cell lysates: cells were lysed and aquired as previously described^101^, using the Orbitrap Fusion Lumos Tribbrid mass spectrometer.

### Mass Spectrometry data Analysis

Raw mass spectrometry files were processed with the MaxQuant computational platform, version 1.6.2.10. Proteins and peptides were identified using the Andromeda search engine by querying the reviewed human Uniprot database (downloaded 09-22-2023 20426 entries). Standard settings with the additional options match between runs, and unique peptides for quantification were selected. The generated ‘proteingroups.txt’ data were imported in R4.30/Rstudio 2024.04.1. ‘reverse’, ‘potential contaminants’ and ‘only identified by site’ peptides were filtered out and label free quantification values were log2 transformed. Missing values were imputed by a normal distribution (width=0.3, shift = 1.8), assuming these proteins were close to the detection limit. Statistical analyses were performed using moderated t-tests in the LIMMA package. A Benjamini-Hochberg adjusted P value <0.05 and absolute log2 fold change >1 was considered statistically significant and relevant.

For supervised classification, we generated theoretical profiles in which conditions of interest intensities were set to an arbitrary high and low in all other samples (Kreft et al. 2023). These profiles were correlated with all statistically significant proteins. Correlation between data and theoretical protein profiles was performed using Pearson correlations, a ρ > 0.5 or 0.6 and Benjamini-Hochberg adjusted P value <0.05 was used as threshold.

Gene ontology overrepresentation analyses were performed using the clusterprofiler package ^102^.

The R package ggplot2 was used for graphical representations.

### Genetic modification of T cells with Cas9 RNPs

crRNAs were designed in Benchling (https://benchling.com; Appendix B). Cas9 RNP production and T cell nucleofection was performed as previously described (Freen-van Heeren et al. 2020). Briefly, Alt-R crRNA and tracrRNA were reconstituted to 100 µM in Nuclease Free Duplex buffer (all IDT). As a negative control, nontargeting negative control crRNA #1 was used (IDT). Oligos were mixed at equimolar ratios (i.e. 4.5 mL total crRNA + 4.5 mL tracrRNA) in nuclease-free PCR tubes and denatured by heating at 95°C for 5 min. Nucleic acids were cooled down to room temperature prior to mixing them with 30 µg Alt-R™ S.p. Cas9 Nuclease V3 (IDT) to produce Cas9 ribonuclear proteins (RNPs). Mixture was incubated at room temperature for at least 10 min prior to nucleofection. For nucleofection, human CD3^+^ T cells activated for 48 h with αCD3/αCD28 were washed twice with PBS and resuspended in P2 buffer (Lonza). Cells were electroporated in 16-well strips in a 4D Nucleofector X unit (Lonza) with program EH100 for human T cells &P2 buffer. Knockout efficiency was determined 5 d after electroporation by Western blot, or by PCR.

### RNA immunoprecipitation

Cytoplasmic lysates of 25×10^6^ human CD3^+^ T cells activated for 3h with αCD3/αCD28 were prepared using lysis buffer (10 mM HEPES, pH 7.0, 100 mM KCl, 5 mM MgCl2, 0.5% NP40) freshly supplemented with 1 mM DTT, 100 U/ml RNase OUT (both Invitrogen), 0.4 mM Ribonucleoside Vanadyl Complex (NEB) and 1% EDTA-free protease/phosphatase inhibitor cocktail (Thermo Scientific). Protein G Dynabeads (Invitrogen) were prepared according to manufacturer’s protocol. The lysate was immunoprecipitated for 4h at 4ºC with 10 µg polyclonal α-UPF1 (ProteinTech) or a polyclonal rabbit IgG isotype control (12–370, Sigma-Aldrich). RNA was extracted directly from beads by using Zymo Quick-RNA Miniprep Kit, and mRNA expression was measured by RT-PCR as described above.

### Quantitative PCR analysis

T cells were activated for indicated time point with αCD3/αCD28. For mRNA half-life measurements, cells were treated for an additional 1h or 2h with 5 µg/mL actinomycin D (ActD) (Sigma-Aldrich). Total RNA was extracted using Quick-RNA MiniPrep plus kip (Zymo). cDNA was synthesized with SuperScript III (Invitrogen), RT-PCR was performed using SYBR Green on a StepOne Plus (Applied Biosystems). Ct values were normalized to RPS18 levels.

### Quantification and statistical analysis

Results are shown as mean ± SD. Statistical analysis was performed with GraphPad Prism 10 with Wilcoxon two-tailed ratio paired or unpaired Student’s t test when comparing two groups, or with one-way ANOVA test with Dunnett correction when comparing more than two groups. p values < 0.05 were considered statistically significant.

Appendix: Key resources

### Appendix A

| Antibodies | Source | Identifier |
| --- | --- | --- |
| Rat anti-mouse IgG2a (clone MW1483) | Sanquin | n/a |
| Mouse anti-CD28 (clone CD28.2) | Biolegend | Cat# 302902; RRID:AB_314304 |
| Mouse anti-CD3 (clone HIT3a) | Biolegend | Cat# 300302; RRID:AB_314038 |
| Pelicluster CD3 | Sanquin | Cat#564364 |
| BUV737 mouse anti-human CD4 (clone SK3) | BD Bioscience | Cat# 564305; RRID:AB_2713927 |
| BUV805 mouse anti-human CD8 (clone SK1) | BD Bioscience | Cat# 612889; RRID:AB_2833078 |
| BUV395 mouse Anti-human CD69 (clone FN50) | BD Bioscience; | Cat# 564364; RRID:AB_2738770 |
| Brilliant Violet 785™ anti-human TNF (clone MAb11) | Biolegend | Cat# 502948; RRID:AB_2565858 |
| eFluor 450 mouse anti-human IFN gamma (clone 4S.B3) | eBiosciences | Cat# 48-7319-42; RRID:AB_2043866 |
| APC anti-human IL2 (clone MQ1-17H12) | Biolegend | Cat# 500310; RRID:AB_3083299 |
| Alexa Fluor® 647 mouse anti-human CD137 (clone 4B4-1) | Biolegend | Cat# 309824; RRID:AB_2566258 |
| LIVE/DEAD™ Fixable Near-IR Dead Cell Stain Kit, for 633 or 635 nm excitation | Invitrogen | Cat# L34976 |
| CellTrace Violet® | Thermo Fisher Scientific |  |
| UPF1 rabbit anti-human Polyclonal antibody | Proteintech | cat# 23379-1-AP ; RRID:AB |
| CSTF2T rabbit anti-human Polyclonal | Proteintech | cat# 12419-1-AP; RRID:AB2230895 |
| ZFP36L1 mouse anti-human monoclonal antibody | Abcam | Cat# ab42473 RRID: AB_883662 |
| ZFP36L2 rabbit anti-mouse polyclonal antibody | Abcam | Cat# ab70775 RRID: AB_1271231 |
| ZFP36 rabbit anti-mouse polyclonal antibody | Origene | Cat# OTI3D10 |
| Mouse anti-RhoGDI | Abnova | Cat# 89-113-917; RRID:AB |
| Rabbit polyclonal IgG | Sigma-Aldrich | Cat# 12-370 |
| Mouse anti-rabbit IgG light chain (HRP) | Abcam | Cat# ab99697; RRID:AB_10673897 |
| **Bacterial and viral strains** |  |  |
| E. coli DH5alpha super competent bacteria |  | Invitrogen Cat#1 8265017 |
| **Biological samples** |  |  |
| Human PBMC | Sanquin | n/a |
| **Chemicals, peptides, and recombinant proteins** |  |  |
| Trypsin Gold, Mass Spectrometry Grade | Promega | Cat# V5280 |
| GolgiPlug Protein Transport Inhibitor (containing Brefeldin A) | BD Biosciences | Cat# BD 555029 |
| Recombinant human IL-15 | Peprotech | Cat# 200-15 |
| Recombinant human IL2 | Peprotech | Cat# 200-02 |
| GeneJammer | Agilent | Cat# 204130 |
| RetroNectin | Takara Bio | Cat# T100B |
| Phorbol 12-myristate 13-acetate (PMA) | Sigma-Aldrich | Cat# P8139 |
| Ionomycin calcium salt | Sigma-Aldrich | Cat# 13909 |
| RNase A | Thermo Scientific | Cat# EN0531 |
| Precision Counting Beads | Biolegend | Cat#424902 |
| GelRed Nucleic Acid Gel Stain | Biotium | Cat# 41003 |
| Actinomycin D | Sigma-Aldrich | Cat# A9415 |
| Alt-R™ S.p. Cas9 Nuclease V3 | DT | ICat# 10000735 |
| Power SYBR™ Green PCR Master Mix | Applied Biosystems | Cat# 4367660 |
| RNase A, DNase and protease-free (10 mg/mL) | Thermo Scientific | Cat# EN0531 |
| RNaseOUT™ Recombinant Ribonuclease Inhibitor | Invitrogen | ICat# 10777019 |
| Ribonucleoside Vanadyl Complex | New England Biolabs | Cat# S1402S |
| Halt™ Protease and Phosphatase Inhibitor Single-Use Cocktail, EDTA-Free (100X) | Thermo Scientific | Cat# 78443 |
| Digitonin, High Purity | Calbiochem | Cat# 300410 |
| Lympholyte-M | Cederlane | Cat# CL5035 |
| Dynabeads™ Protein A for Immunoprecipitation | Invitrogen | Cat# 10008D |
| Dynabeads™ Protein G for Immunoprecipitation | Invitrogen | Cat# 10003D |
| Streptavidin Sepharose High Performance | GE Healthcare | Cat# GE17-5113-01 |
| **Critical commercial assays** |  |  |
| NucleoSpin Plasmid Transfection grade, Mini kit for ultrapure plasmid DNA | Machery-Nagel |  |
| Fixation/Permeabilization Solution Kit with BD GolgiPlug™ | BD Biosciences | Cat# 555028 |
| P2 Primary Cell 4D-Nucleofector™ X Kit S | Lonza | Cat# V4XP-2032 |
| SuperScript™ III First-Strand Synthesis System | Invitrogen | Cat# 18080051 |
| **Equipment** |  |  |
| VerityPro Thermal Cycler | Thermo Scientific | n/a |
| Mupid One Electrophoresis System | Nippon Genetics Europe | n/a |
| NGS platform | Illumina | n/a |
| NanoDrop 2000 | Thermo Scientific | n/a |
| BD LSR Fortessa Cell Analyzer | BD Biosciences | n/a |
| FACS Symphony A5 Cell Analyzer | BD Biosciences | n/a |
| Amersham ImageQuant 800 | Cytiva | n/a |
| Casy Cell counter | Roche Innovatis | n/a |
| 4D-Nucleofector® X Unit | Lonza | Cat# AAF-1003X |
| **Deposited data** |  |  |
| Raw data files Mass Spectrometry analysis (ProteomeXchange) | This paper | PRIDE: PXD056415 |
| **Experimental models: Cell lines** |  |  |
| Human: FLYRD18 cells | ECACC | Cat# 95091902 |
| Human: Jurkat cell line |  |  |
| **Recombinant DNA** |  |  |
| GFP-TIS11B | <sup>28</sup> | Addgene Cat # 121157 |
| pSF3-UltraID | <sup>30</sup> | Addgene Cat # 172878 |
| **Software and algorithms** |  |  |
| FlowJo version 10.8.1 | BD Biosciences | https://www.flowjo.com/ |
| GraphPad Prism version 9.1.1 | GraphPad Software | https://www.graphpad.com/ |
| R environment for statistical computing | The R foundation | https://www.r-project.org/ |
| RStudio Version 2024.12.1+563 | RStudio | https://www.rstudio.com/ |
| StepOne Software v2.3 | Applied Biosystems | https://www.thermofisher.com/nl/en/home/technical-resources/softwaredownloads/StepOne-andStepOnePlus-Real-Time-PCRSYSTEM.html |
| Maxquant 1.6.10 | <sup>103</sup> | https://www.maxquant.org/ |

### Appendix B

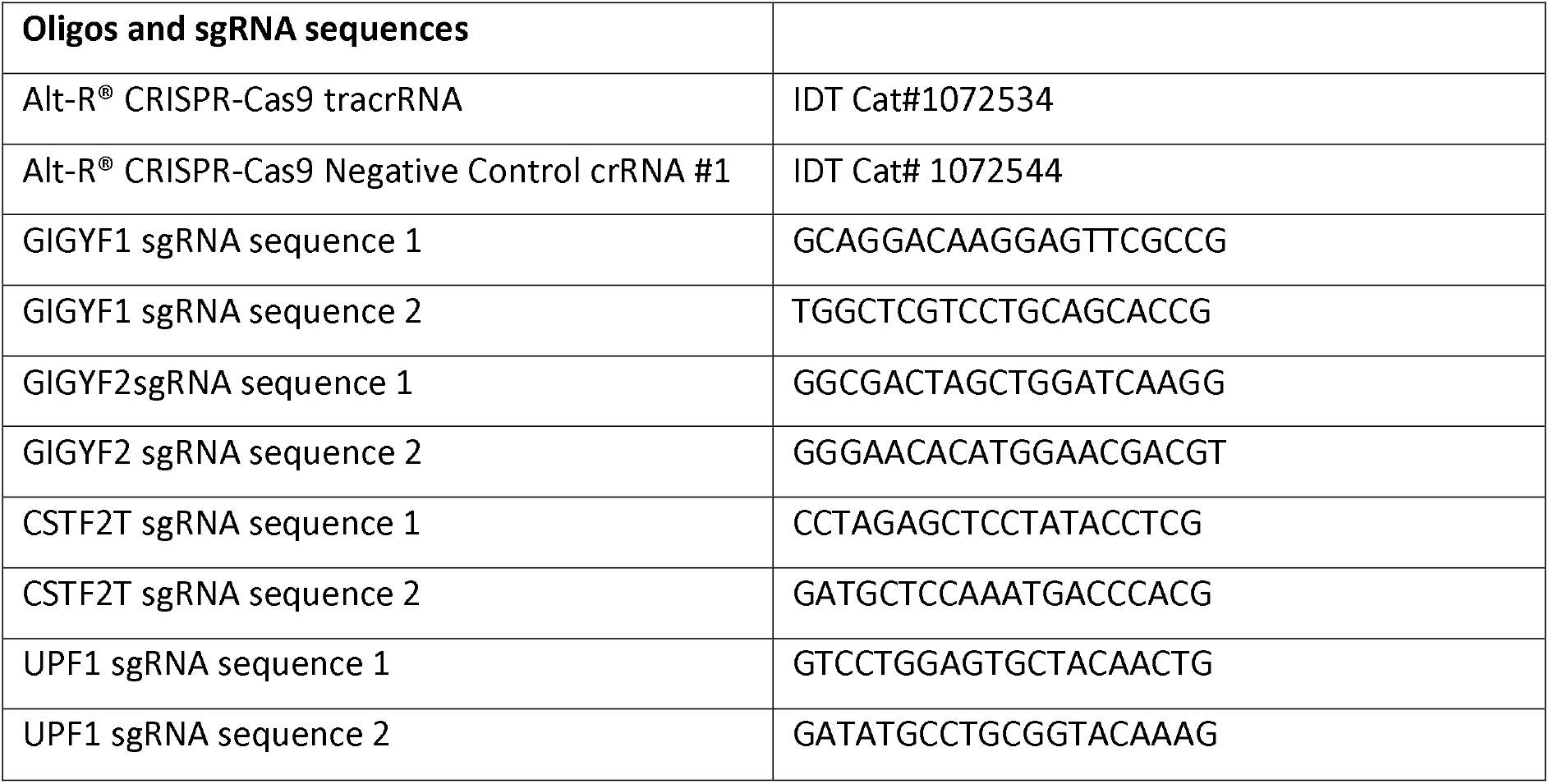

## SUPPLEMENTARY FIGURE LEGENDS

**Supplemental Figure 1.**
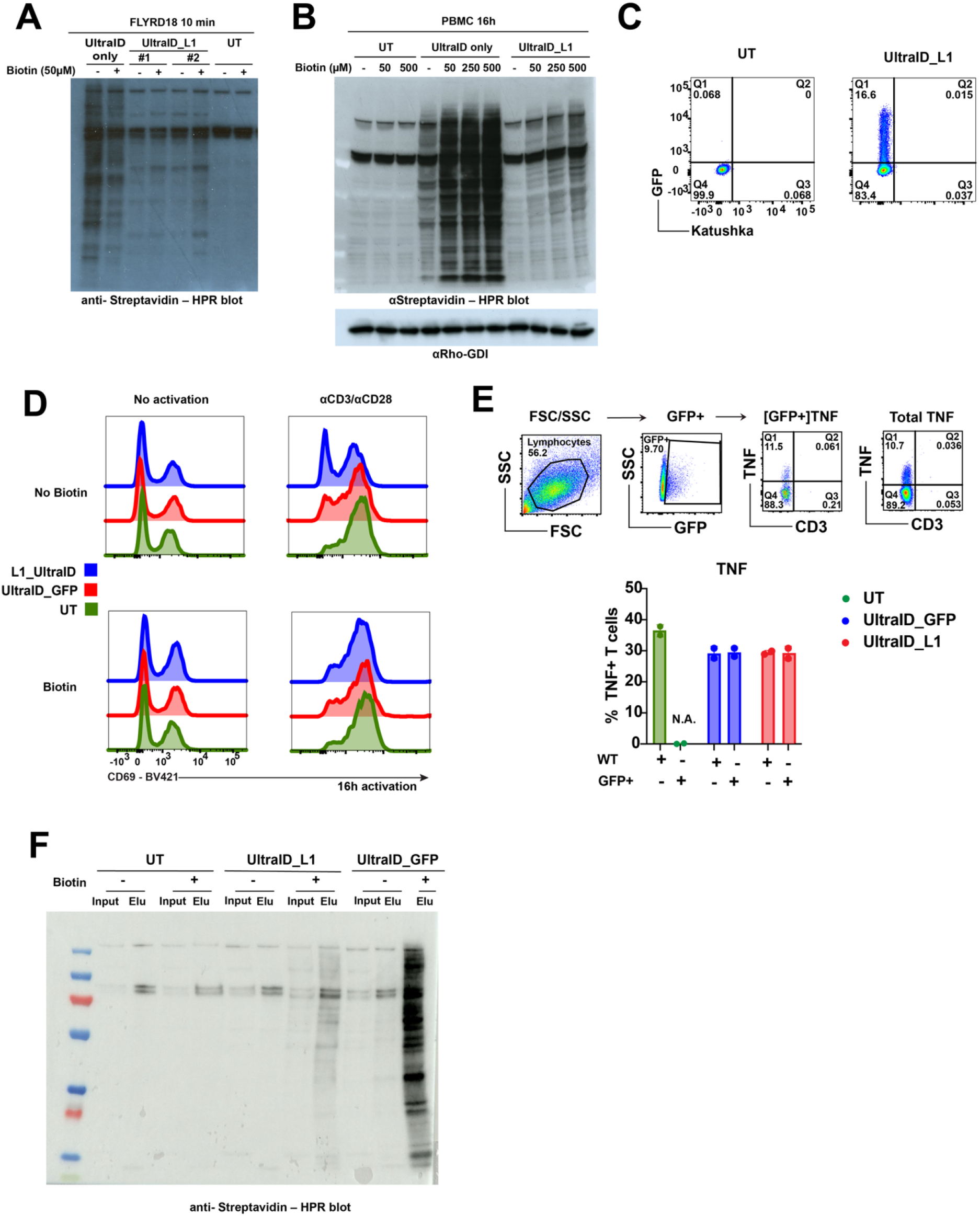
Optimizing Proximity labeling in primary human T cells. **a**. Immunoblot of biotinylated proteins with aStreptavidin-HPR of FLYRD18 cells left untransduced (UT) or transduced with indicated constructs (UltraID_L1 with two different DNA preparations and UltraID_only) in the presence or absence of ±50µM biotin for 10 min. **b**. Immunoblot of T cells transduced with UT, UltraID_L1 or UltraID_only in the presence or absence of aCD3/aCD28 and with indicated concentrations of Biotin for 16h. Rho-GDI was used as loading control. Representative of at 6 donors in two independently performed experiments. **c**. Flow cytometry plot of transduction efficiency based on GFP expression. **d**. CD69 expression of untransduced (UT), UltraID_L1 (Blue) or UltraID_GFP (Red) T cells that were rested for 7 days, or that were re-activated with aCD3/aCD28 for 16h in the presence or absence of 500µM Biotin. **e**. UT (Green), UltraID_GFP (Red), and UltraID_L1 (Blue) transduced T cells were reactivated with aCD3/aCD28 for 5h. Top: gating strategy. Bottom: percentage of TNF expressing T cells. **f**. Enrichment of biotinylated proteins as visualized by Streptavidin blot of 10% input and 10% of the final elution of T cells transduced with UltraID_L1 or UltraID_GFP, and that were re-activated with aCD3/aCD28 for 16h in the presence of biotin. Representative of at least 2 independent experiments with 3 donor pools. *See Figure 1*

**Supplemental Figure 2.**
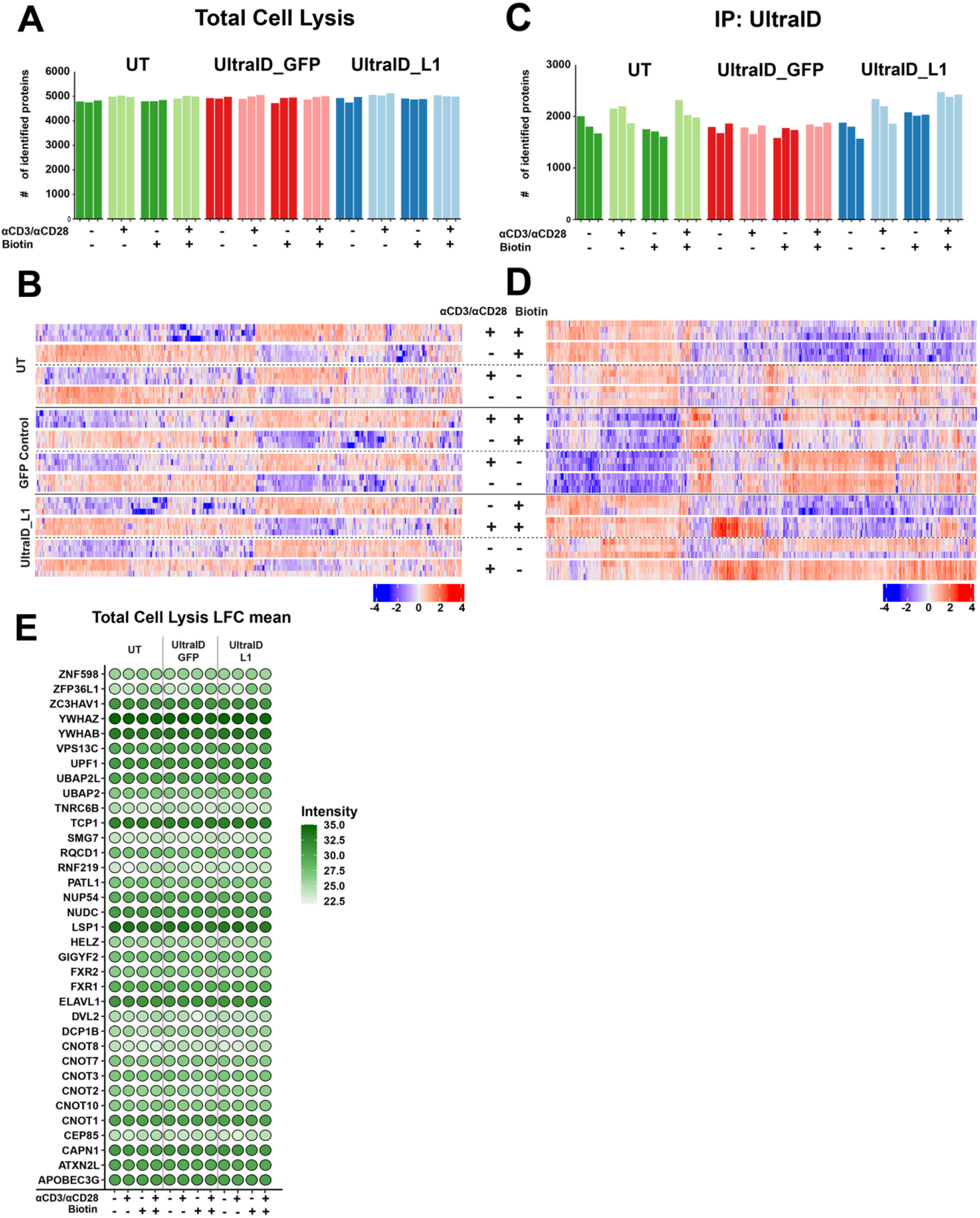
Proximity labeling identifies the ZFP36L1 interactome in primary human T cells. **a**. Number of identified proteins of the total Cell Lysis (TCL) by Mass Spectrometry (MS). Each bar represents one donor pool of 40 donors. **b**. Heatmap of all proteins that were significantly enriched in one of the 12 conditions in Total Cell Lysate (5250 proteins, LFC >1, p.adj<0.05). Color scale represents Z-scored log2 median-centered averaged intensities. **c**. Number of identified proteins after UltraID_L1 streptavidin pulldown (IP) by MS. Each bar represents one donor pool of 40 donors. **d**. Heatmap of proteins that were significantly enriched in the UltraID-L1 IP: T cell screen 1 (2439 proteins, LFC >1, p.adj<0.05). Color scale represents Z-scored log2 mediancentered averaged intensities. **e**. LFC mean intensities of protein expression in the total cell lysates that were found to interact with ZFP36L1 as depicted in Figure 2B. Proteins with LFC >1, p.adj<0.05 are considered significantly enriched. Scale represents LFC average intensities. Mean of 3 donor pools. *See Figure 2*

**Supplemental Figure 3.**
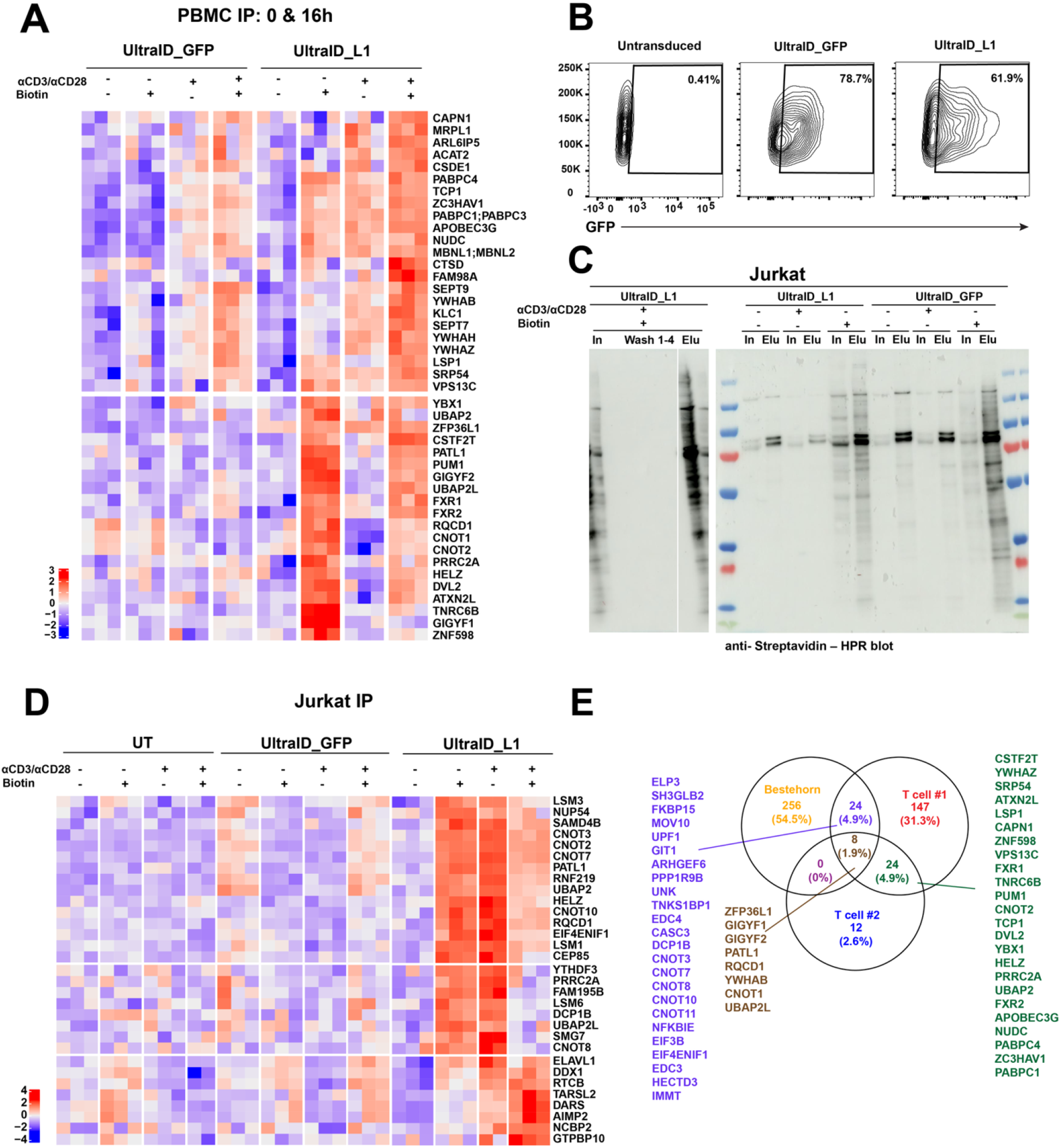
Proximity labeling identifies the ZFP36L1 interactome in Jurkat cells. **a**. Repeat experiment of UltraID Pull down for ZFP36L1 (T cell screen 2). Heatmap of supervised classification displaying biotinylated proteins that were significantly enriched. Correlation between hits and disorder-specific theoretical protein profiles had a Pearson correlation of >0.5, p.adj <0.05, LFC>1. Color scale represents Z-scored log2 mediancentered averaged intensities. **b**. Flow cytometry analysis of GFP-expressing Jurkat cells transduced with indicated constructs. Representative of 3 technical replicates. **c**. Streptavidin immunoblot of input (10%), flow through and consecutive washes (1-4) of the pulldown, in addition to the elution (10%). Representative of at least 2 independent experiments with 3 technical replicates. **d**. Heatmap of supervised classification displaying biotinylated proteins that were enriched in Jurkat UltraID_L1 16h screen. Correlation between hits and disorder-specific theoretical protein profiles had a Pearson correlation of >0.5 and p.adj <0.05 and LFC>1. Color scale represents Z-scored log2 median-centered averaged intensities. **e**. Venn diagram of ZFP36L1 protein interactions from the UltraID T cell screen 1 and 2 identified in FIg xxx, compared with the ZFP36L1 interactome screen in HEK 293T cells from^42^ *See Figure 2*

**Supplemental Figure 4.**
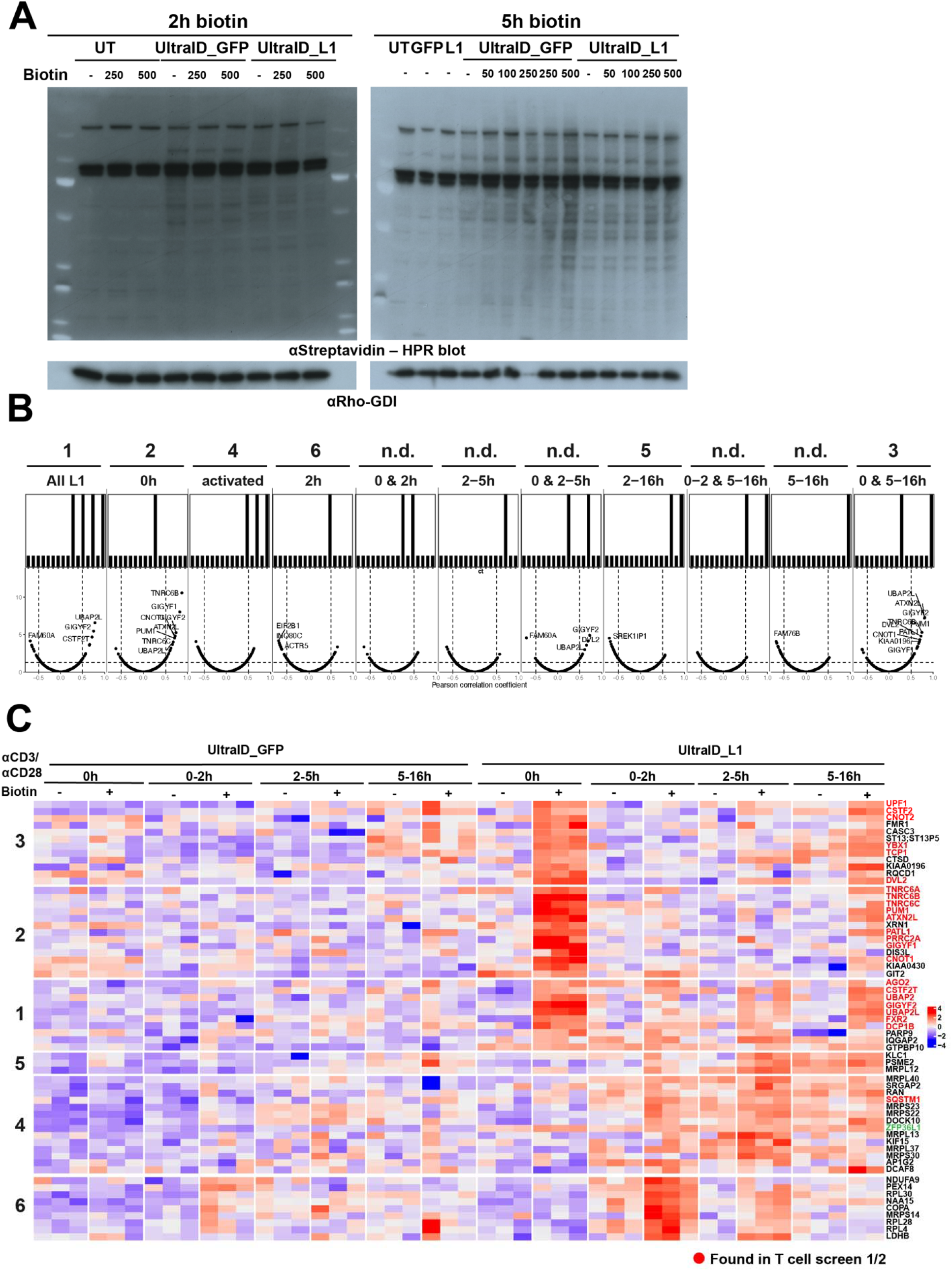
Time-resolved proximity labeling reveals dynamic interactions with ZFP36L1. **a**. Immunoblot of biotinylated proteins of indicated T cells after 2h and 5h of activation with aCD3/aCD28. Indicated concentration of biotin was added during the entire time of T cell activation. Rho-GDI served as loading control. Blot representative of 3 independent donors. **b**. Shape analysis of biotinylated proteins from the T cell activation time course (Figure 5B) using supervised classification, identified to contain interaction partners with a Pearson correlation of >0.5, and p.adj <0.05 and LFC>1. Mean of 3 donor pools. **c**. Heatmap of supervised classification of biotinylated proteins enriched in the shape analysis from panel b. Proteins marked in red were also identified in T cell screen 1 and/or 2. Color scale represents Z-scored log2 median-centered averaged intensities. *See Figure 4*

**Supplemental Figure 5.**
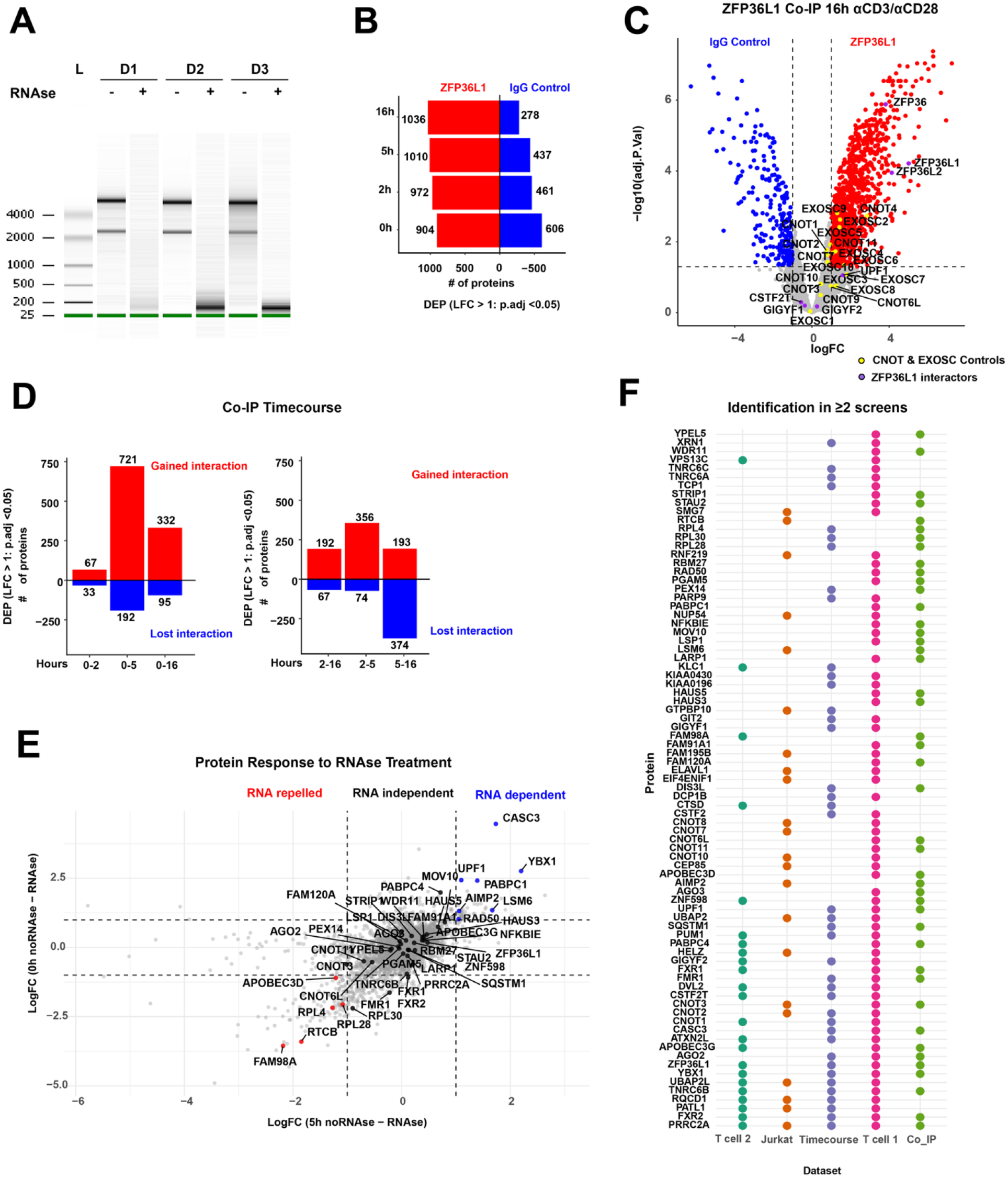
Co-IP reveals RNA-dependent and -independent interactions of ZFP36L1. **a**. RNA profile of T cell lysates measured by Bioanalyser trace with or without treatment throughout entire protocol with RNAse A (200 U/mL) that were used for the ZFP36L1 Co-IP. (L) = ladder. N= 3 donor pools. **b**. Number of proteins that were significantly enriched in the ZFP36L1 Co-IP (Red) vs IgG controls (Blue) (LFC >1, p.adj <0.05) at different time points of T cell activation. **c**. Volcano plot of proteins identified in the ZFP36L1 Co-IP at 16h of T cell activation. Only proteins identified in all three replicates were considered putative interactors. Yellow dots indicate CNOT and EXOSC proteins. Purple dots indicate proteins that were among the top ZFP36L1 interactors in the UltraID_L1 interactome screens. **d**. umber of proteins identified as gained (red) or lost (blue) in ZFP36L1 Co-IP upon RNAse A treatment at indicated time points, compared to the IgG control. Proteins with p.adj <0.05 and LFC>1 or LFC >-1 were considered significant. **e**. Scatter plot analysis of protein responses to RNase treatment between 0h and 5h of αCD3/αCD28 stimulation. Proteins are classified as RNA-repelled (red), RNA-independent (black), or RNA-dependent (blue). Significance threshold of >1 LFC is indicated by dashed lines. **f**. Dot plot depicting protein identification across multiple datasets (T cell 1& 2, Jurkat, Timecourse, Co_IP). Each row represents a different protein, with colored dots indicating presence/detection in each dataset. Proteins are only shown when identified in >2 screens. *See Figure 5*

**Supplemental Figure 6.**
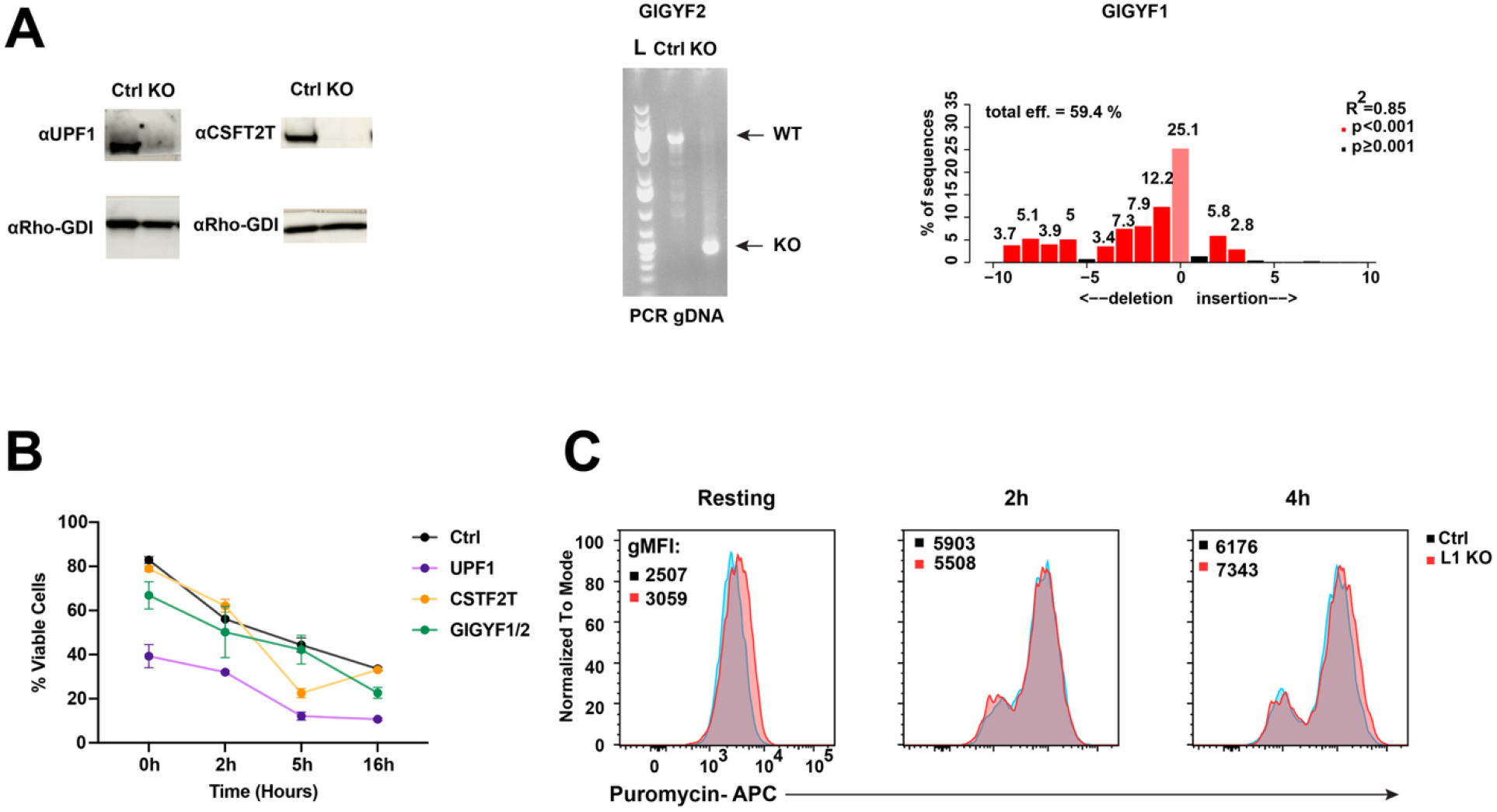
UPF1 and GIGYF1/2 deletion alters T cell viability. **a**. Validation of CRISPR-Cas9 gene-editing for indicated RBPs. 6 days after gene editing, efficiency was determined by immunoblot (left panel: UPF1 and CSFT2T, with Rho-GDI as loading control), by PCR on genomic DNA (middle panel: GIGYF2; arrows indicate PCR product of WT and KO product), or by TIDE analysis (right panel: GIGYF1 KO cells). Representative of n=9 independent donors. **b**. Cell viability of indicated T cells as defined by IR-dye through flow cytometry at indicated time points of activation post αCD3/αCD28 re-stimulation. n= 2 donors. **c**. Puromycin incorporation assay in ZFP36L1 KO versus WT T cells activated with PMA/Iono for indicated time points, as quantified by flow cytometry. Numbers indicate gMFI values. Representative of 3 donors. *See Figure 6*

## Acknowledgements

We thank A. Guislain and S. Toll for technical. help and N. Servaas and K. Rooijers for critical reading of the manuscript. This research was supported by Oncode Institute, the European Research Council consolidator award PRINTERS 817533, and Landsteiner Foundation for Blood Transfusion Research grant 2202, all to M.C.W.; and the Landsteiner Foundation for Blood Transfusion Research grant 2103 to B.P. and M.C.W.

## AUTHOR CONTRIBUTIONS

Conceptualization, A.P.J. and M.C.W.; Methodology, A.P.J., B.P., M.v.B., J.B. and M.C.W.; Investigation, A.P.J., L.W., A.B., F.P.J.v.A., C.v.d.Z.; Formal Analysis, A.P.J., and A.J.H..; Validation, A.P.J., S.E., and A.B.; Writing-Original Draft, A.P.J and M.C.W.; Writing – Review & Editing, A.P.J., B.P., A.J.H., F.P.J.v.A., C.v.d.Z., M.v.B., J.B., and M.C.W.; Supervision, M.C.W.; Funding Acquisition, M.C.W.

## MATERIALS & CORRESPONDENCE

M.C. Wolkers

## MATERIALS AVAILABILITY

This study did not generate new unique reagents.

## DATA CODE AND AVAILABILITY

- The mass spectrometry proteomics data of the RNA Aptamer Pulldown have been deposited to the ProteomeXchange Consortium via the PRIDE^104^ partner repository with the dataset identifier PXD061869.
- This paper does not report original custom code. All codes used in this paper are available from the lead contact upon request.
- Any additional information required to reanalyze the data reported in this paper is available from the lead contact upon request.

